# Bioinformatics analyses of significant genes, related pathways and candidate prognostic biomarkers in Alzheimer’s disease

**DOI:** 10.1101/2021.05.06.442918

**Authors:** Basavaraj Vastrad, Chanabasayya Vastrad

## Abstract

Alzheimer’s disease (AD) is one of the most common causes of dementia and frailty. This study aimed to use bioinformatics analysis to identify differentially expressed genes (DEGs) in AD. The Expression profiling by high throughput sequencing dataset GSE125583 was downloaded from the Gene Expression Omnibus (GEO) database and DEGs were identified. After assessment of Gene Ontology (GO) terms and pathway enrichment for DEGs, a protein–protein interaction (PPI) network, module analysis, miRNA-hub gene regulatory network construction and TF-hub gene regulatory network were conducted via comprehensive target prediction and network analyses. Finally, we validated hub genes by receiver operating characteristic curve (ROC). In total, 956 DEGs were identified in the AD samples, including 479 up regulated genes and 477 down regulated genes. Functional enrichment analysis showed that these DEGs are mainly involved in the neuronal system, GPCR ligand binding, regulation of biological quality and cell communication. The hub genes of PAK1, ELAVL2, NSF, HTR2C, TERT, UBD, MKI67, HSPB1, PYHIN1 and TES might be associated with AD. The diagnostic value and expression levels of these hub genes in AD were further confirmed by ROC analysis. In conclusion, we identified pathways and crucial candidate genes that affect the outcomes of patients with AD, and these genes might serve as potential therapeutic targets.

## Introduction

Alzheimer’s disease (AD) is a complex, dementia that mainly affects the memory, cognitive, and behavior functions [1]. Approximately over 45 million elderly people worldwide are affected with AD worldwide in 2020 which is expected to rise to 115 million by the year 2050 [2]. AD involves the degeneration of nervous tissues [3]. Numerous factors might affect AD progression, including genetic (70%) and environmental (30%) [4]; however, the cause and potential molecular mechanism of AD are still unclear. Therefore, there is a crucial commitment to establish new diagnostic strategies and therapeutic agents to develop the prognosis of patients with AD.

To find out novel genes and signaling pathways that is linked with AD and patient prognosis, which might not only help to elucidate the underlying molecular mechanisms involved, but also to discover new biomarkers and therapeutic targets. Apolipoprotein E epsilon 4 (APOE) [5], amyloid-β precursor protein (APP) [6], OPRM1 and OPRL1 [7], RAB10 [8], IL6 [9], Wnt/beta-catenin signaling pathway [10], glycogen synthase kinase-3 signaling pathway [11], insulin signaling pathway [12], CREB signaling pathway [13] and autotaxin-lysophosphatidic acid signaling pathway [14] are linked with development of AD. Therefore, identifying genes and the signal pathways, is essential for AD.

With the rapid advancement of bioinformatics such as RNA sequencing, some high throughput platforms for analysis of gene expression are frequently used to find the differentially expressed genes (DEGs) during disease progression. Now, through gene expression profiling studies using RNA sequencing, more and more DEGs linked with AD have been found.

Therefore, in the present study, we downloaded the original data (GSE125583), provided by Srinivasan et al [15], from the publically available Gene Expression Omnibus database (GEO, http://www.ncbi.nlm.nih.gov/geo/) [16] to identify DEGs and the associated biological processes between AD and healthy control using comprehensive bioinformatics analyses. The DEGs were subjected to gene ontology (GO) enrichment and pathway analyses; moreover, a protein-protein interaction (PPI) network, modules, miRNA-hub gene regulatory network construction and TF-hub gene regulatory network construction were constructed and analyzed to screen for hub genes, miRNAs and TFs. Finely, hub genes were validated by receiver operating characteristic curve (ROC). The aim of this investigation was to find out hub genes and pathways in AD using bioinformatics methods and to then study the potential molecular mechanisms of AD.

## Materials and methods

### Data resources

Expression profiling by high throughput sequencing dataset GSE125583 [15] was downloaded from the GEO database. The data was produced using a GPL16791 Illumina HiSeq 2500 (Homo sapiens). The GSE125583 dataset contained data from 289 samples, including 219 AD samples and 70 normal control samples.

### Identification of DEGs

The identification of DEGs between AD and normal control was performed in limma package in R software [17], an tool designed to compare different groups of samples. The P values were adjusted to correct for the occurrence of false positive results by using the Benjamini and Hochberg False Discovery Rate method [18]. An adjusted P□<□.05, and a |log_2_FC| > 0.731 for up regulated genes and |log_2_FC| < −0.696 for down regulated genes were used as the cutoff values for identifying DEGs. Volcano plots of DEGs were provided using gplot package in R. The ggplot2 package in R was used to perform heatmap of up and down regulated genes in AD and healthy control.

### GO and REACTOME pathway enrichment analysis of DEGs

GO (http://geneontology.org/) [19] and REACTOME (https://reactome.org/) [20] pathway enrichment analyses were executed using g:Profiler (http://biit.cs.ut.ee/gprofiler/) [21]. All DEG symbols were input as a list into the g:Profiler website. The results of GO and REACTOME pathway enrichment analyses were provided by bioinformatics tools in the website. There are processes in GO analysis involved biological processes (BP), cellular components (CC) and molecular functions (MF). The pathways of REACTOME involved metabolism, genetic information processing, environment information-related processes, cell physiological processes and drug research.

### PPI network establishment and modules selection

Protein–protein interaction (PPI) analysis was performed using the Human Integrated Protein-Protein Interaction rEference (HIPPIE) interactome database (http://cbdm-01.zdv.uni-mainz.de/~mschaefer/hippie/network.php) [22], which is software used to explore interactions among proteins. All the symbols of DEGs were inputted. The network file among proteins was downloaded. Cytoscape software (version 3.8.2) (www.cytoscape.org) [23] is an excellent application for network visualization of proteins and was used to generate the PPI network plot constructed by nodes and edges. The hub genes were predicted by Network Analyzer, which is a plugin of Cytoscape to screen hub genes from numerous candidates. The ranks of the hub genes in the network were determined by the value of node degree [24], betweenness centrality [25], stress centrality [26] and closeness centrality [27], which are an algorithm provided by Network Analyzer. Because the node degree, betweenness centrality, stress centrality and closeness centrality values of genes are directly related to its importance, genes with a high node degree, betweenness centrality, stress centrality and closeness centrality value are more likely to be hub genes. The PEWCC1 (http://apps.cytoscape.org/apps/PEWCC1) [28], a plugin for Cytoscape, was used to screen the modules of the PPI network. The criteria were set as follows: degree cutoff=2, node score cutoff=0.2, k-core=2 and maximum depth=100. Moreover, the function and pathway enrichment analysis were performed for DEGs in the modules.

### MiRNA-hub gene regulatory network construction

miRNet database (https://www.mirnet.ca/) is a web-based tool developed by Fan and Xia [29] that supplies the largest available collection of predicted and experimentally verified miRNA-hub gene interactions with various novel and unique features. In the current investigation, hub genes of the DEGs were identified using miRNet database. Furthermore, Cytoscape (version 3.8.2) software [23] was used to establish the miRNA-hub gene regulatory network.

### TF-hub gene regulatory network construction

NetworkAnalyst database (https://www.networkanalyst.ca/) is a web-based tool developed by Zhou et al [30] that supplies the largest available collection of predicted and experimentally verified TF-hub gene interactions with various novel and unique features. In the current investigation, hub genes of the DEGs were identified using NetworkAnalyst database. Furthermore, Cytoscape (version 3.8.2) software [23] was used to establish the miRNA-hub gene regulatory network.

### Validation of hub genes by receiver operating characteristic curve (ROC) analysis

The ROC curves were used to explore the sensitivity and specificity of hub genes for AD diagnosis using R packages “pROC” [31].To further assess the predictive accuracy of the hub genes, ROC analysis was performed to discriminate AD from normal. The area under the ROC curve (AUC) was calculated and used to compare the diagnostic value of these genes.

## Results

### Identification of DEGs

Via limma package in R software, 956 DEGs (479 up regulated genes and 477 down regulated genes) (Table 1) in GSE125583 were extracted after gene expression profile data processing and standardization with the cutoff standard of adjusted P□<□.05, and a |log_2_FC| > 0.731 for up regulated genes and |log_2_FC| < - 0.696. The volcano plot of all DEGs in dataset was shown in Fig. 1. The DEGs were clustered in the heatmap between patients with AD patients and healthy control (Fig. 2).

**Table 1.**
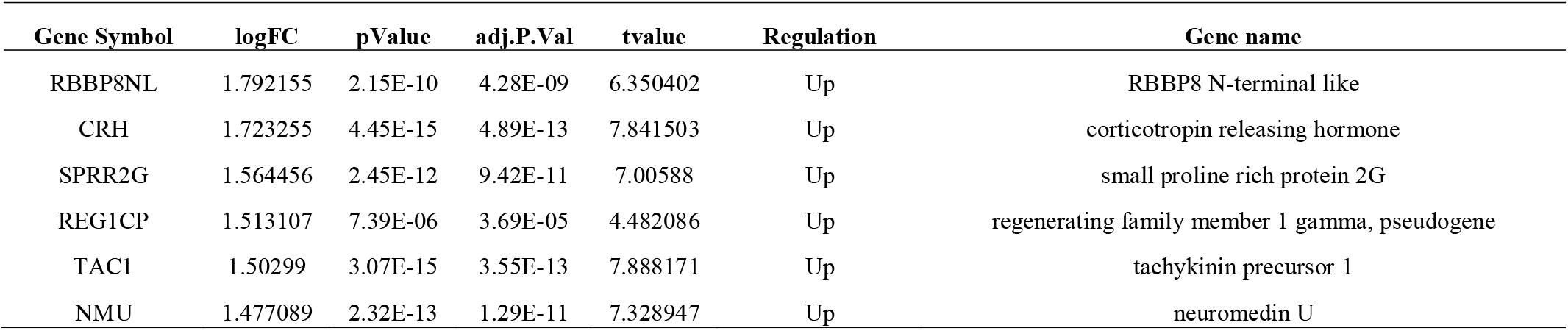
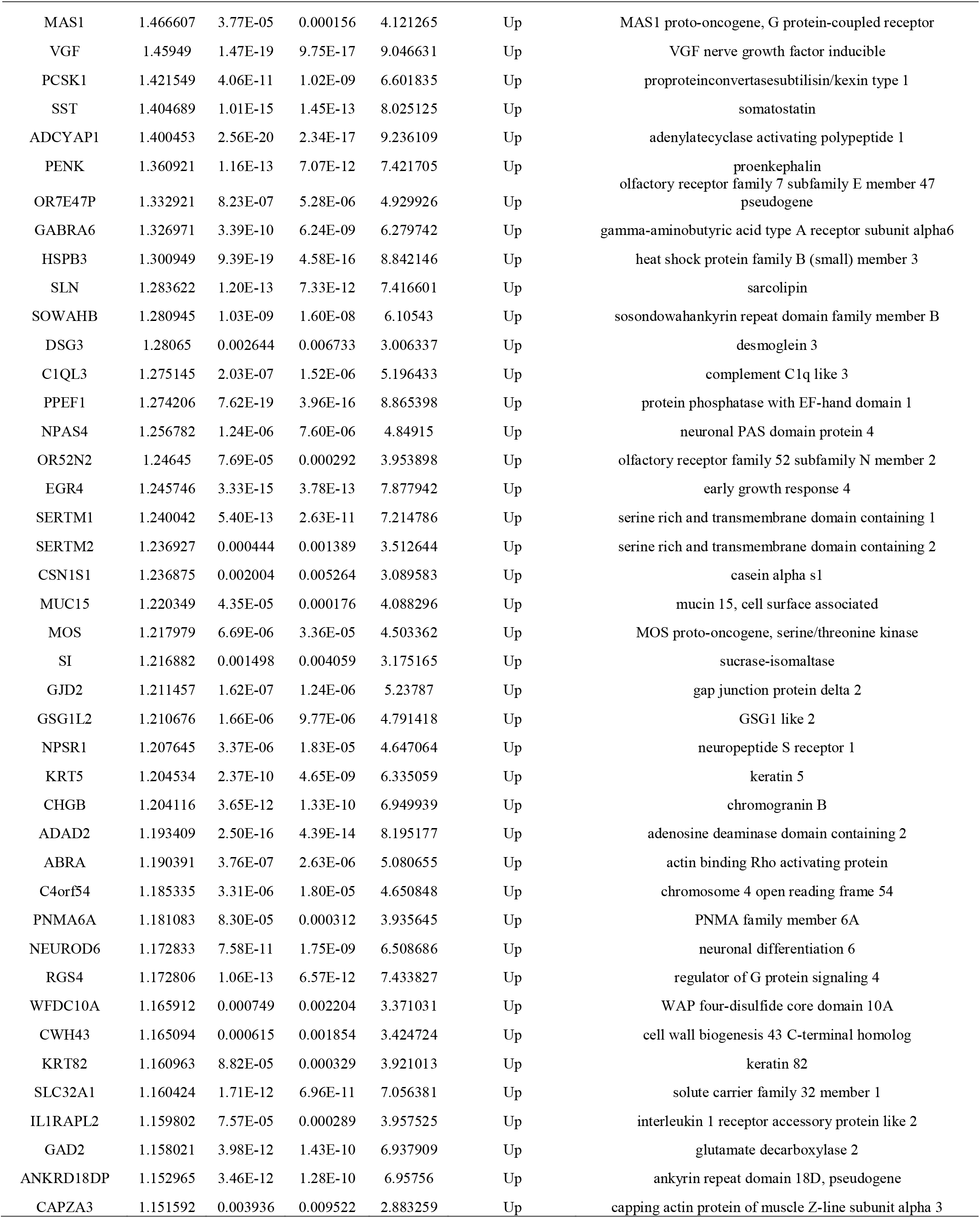
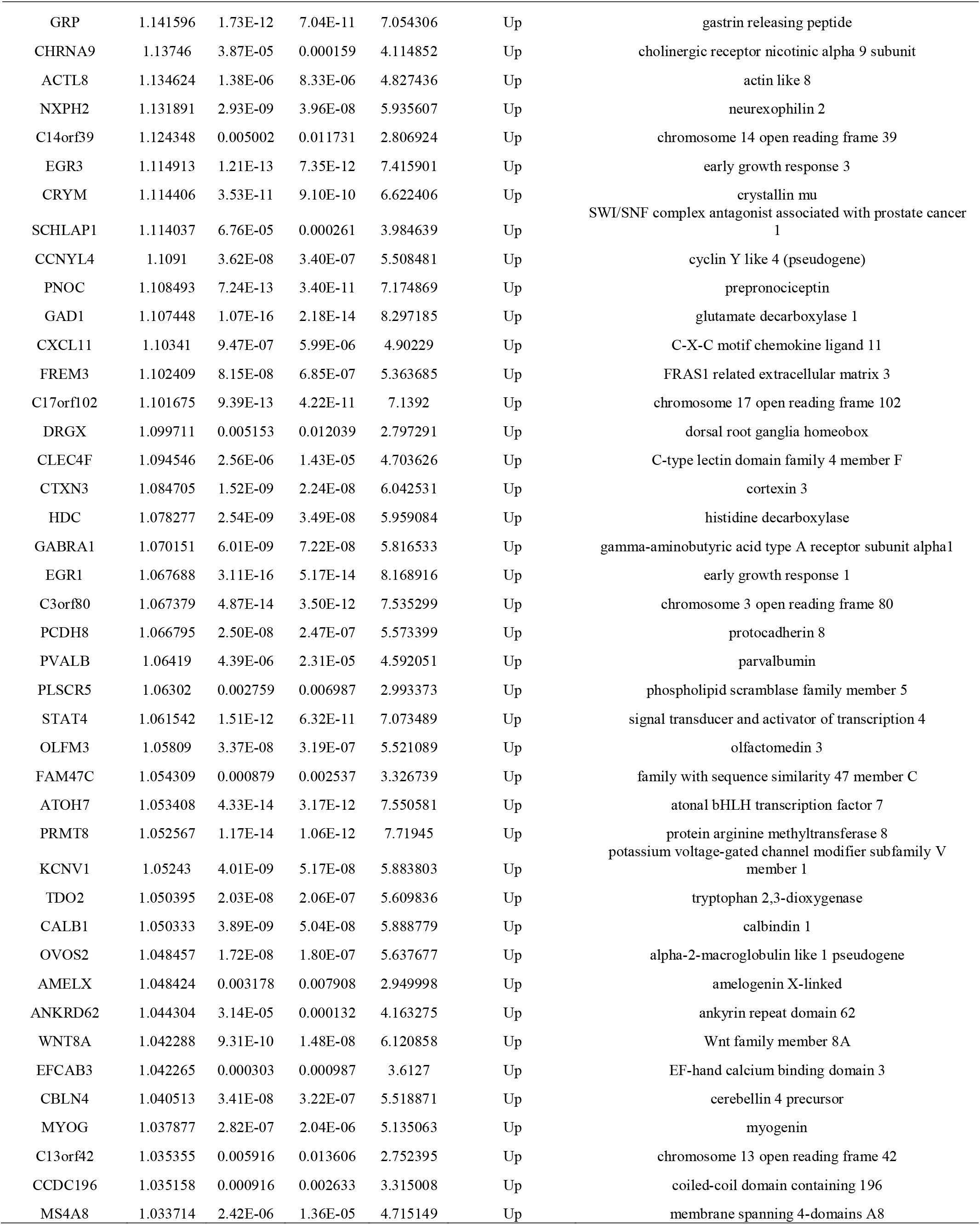
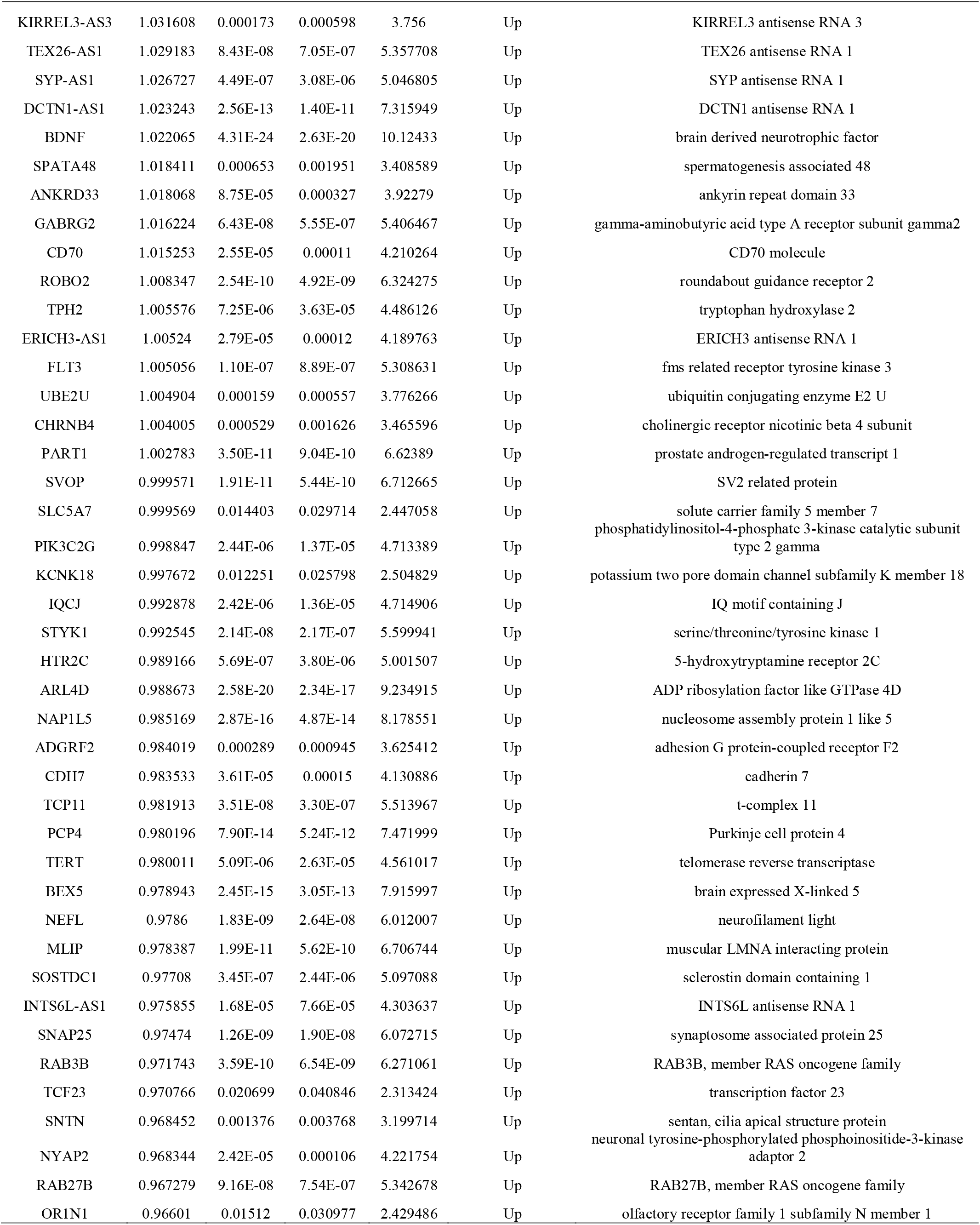
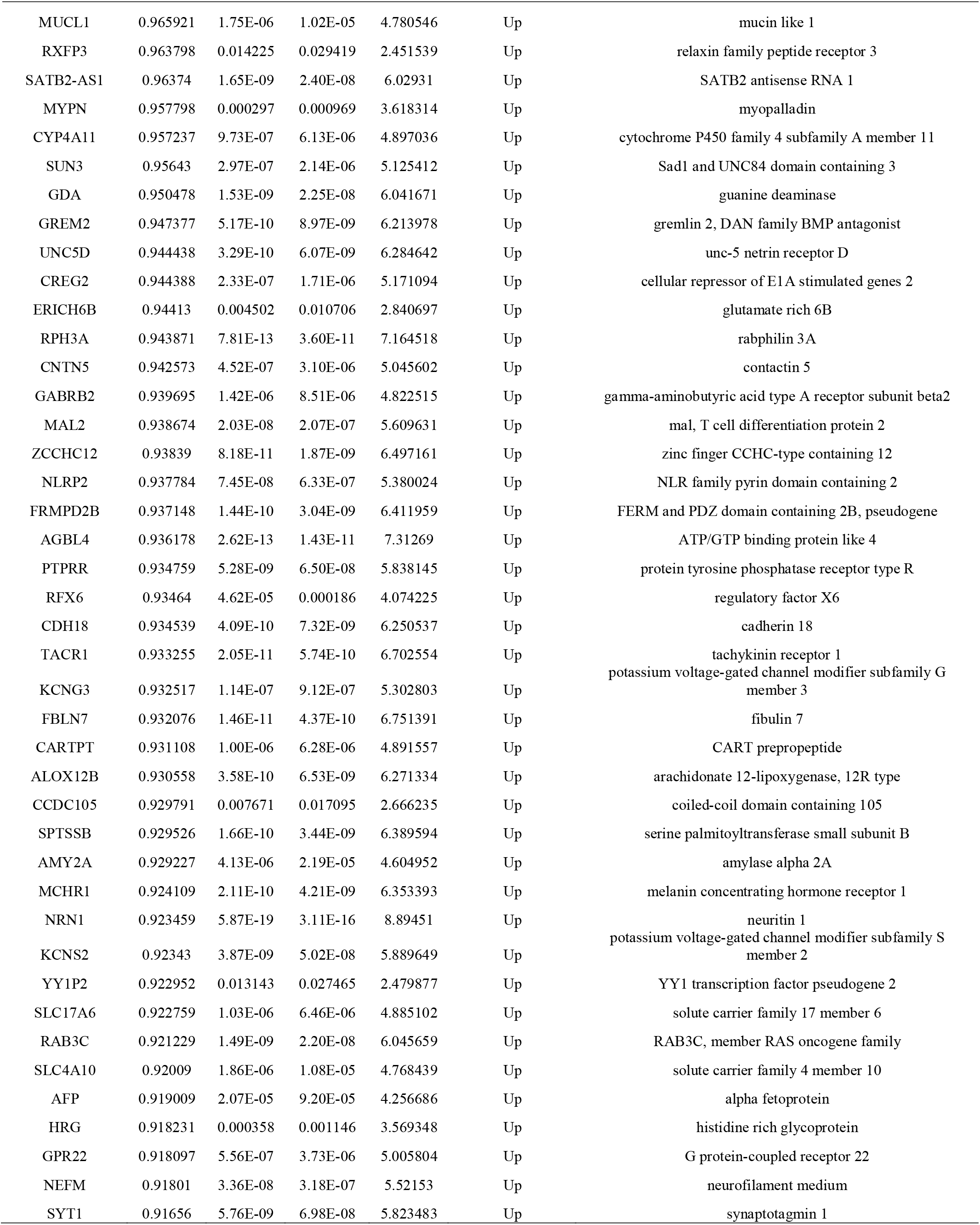
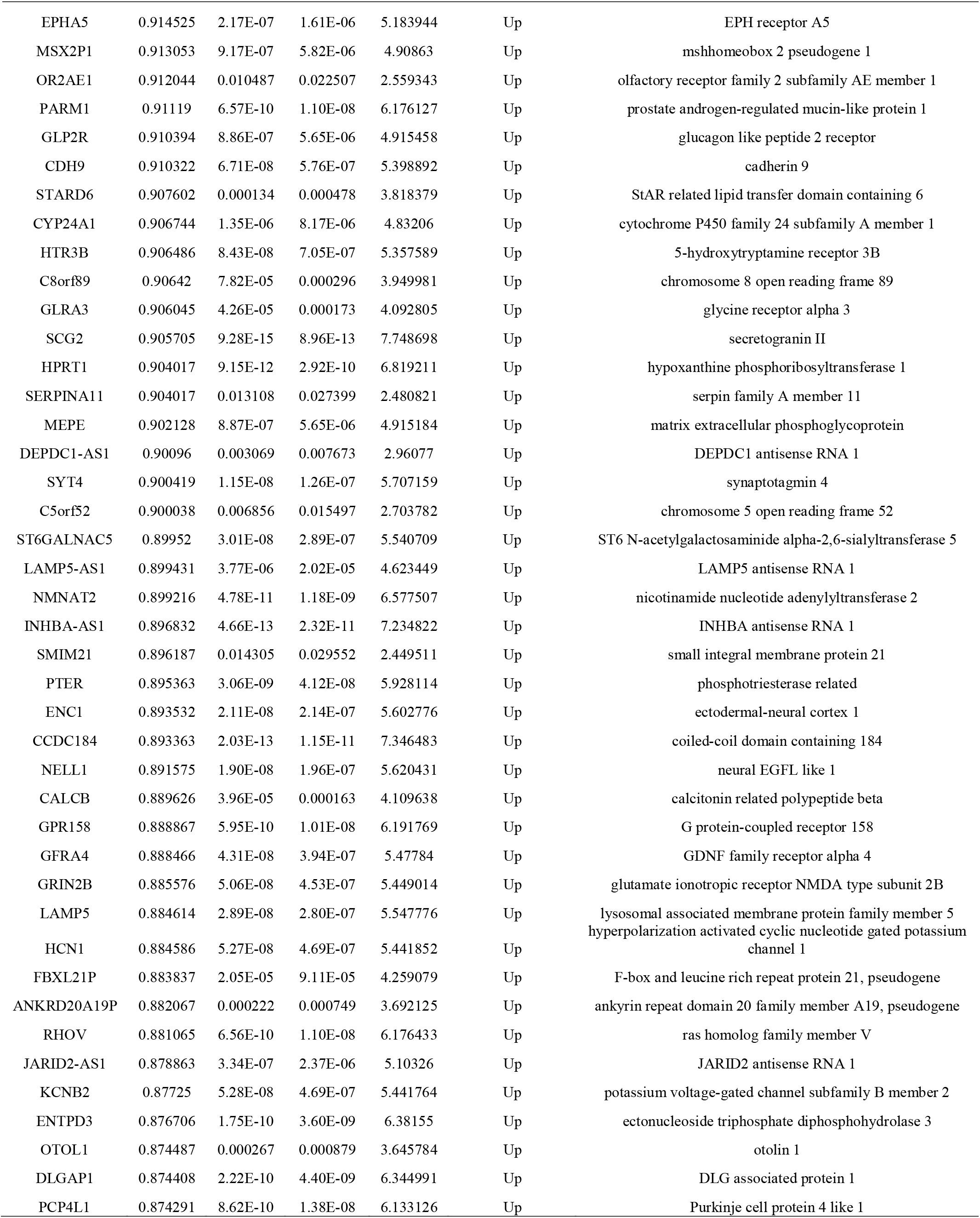
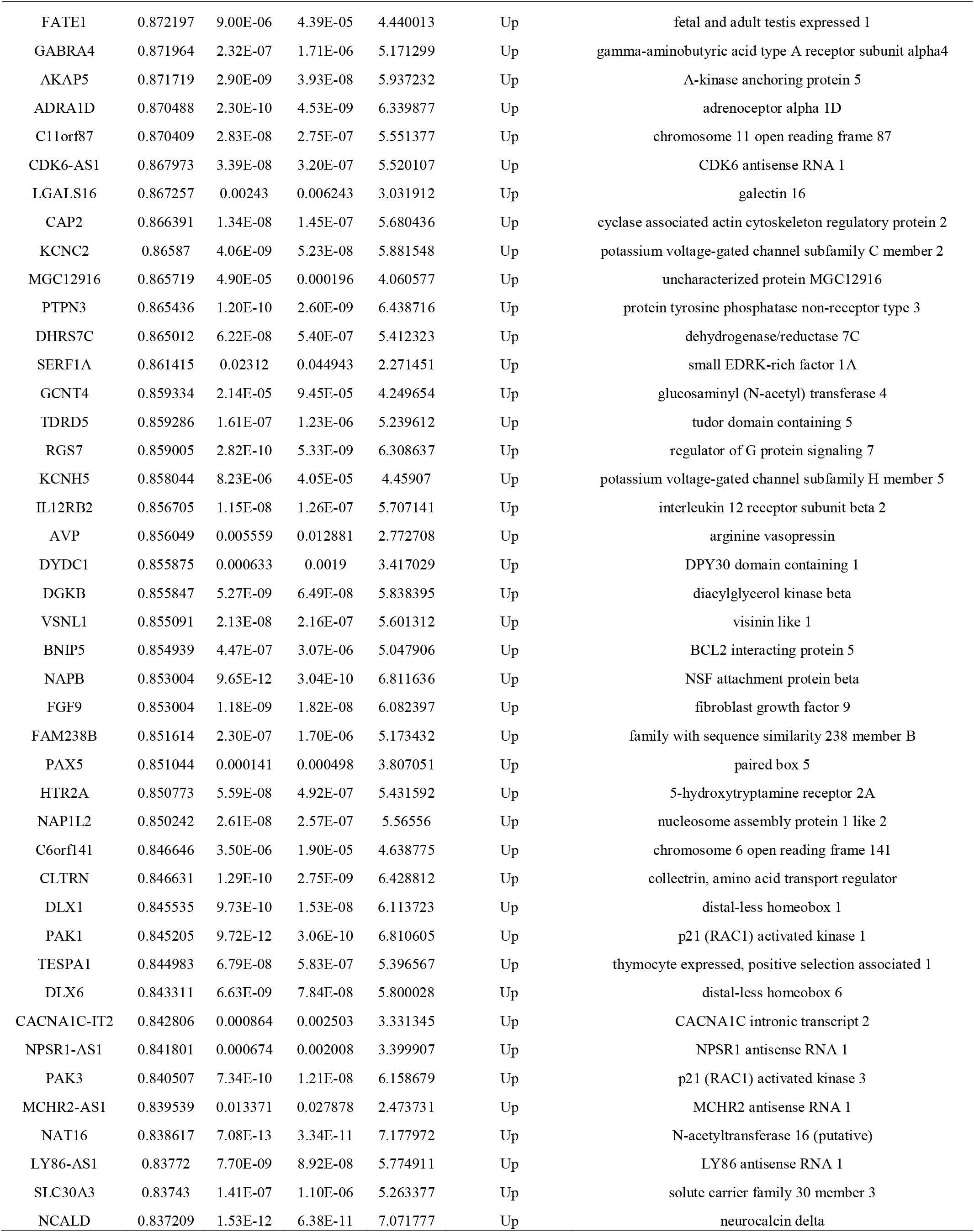
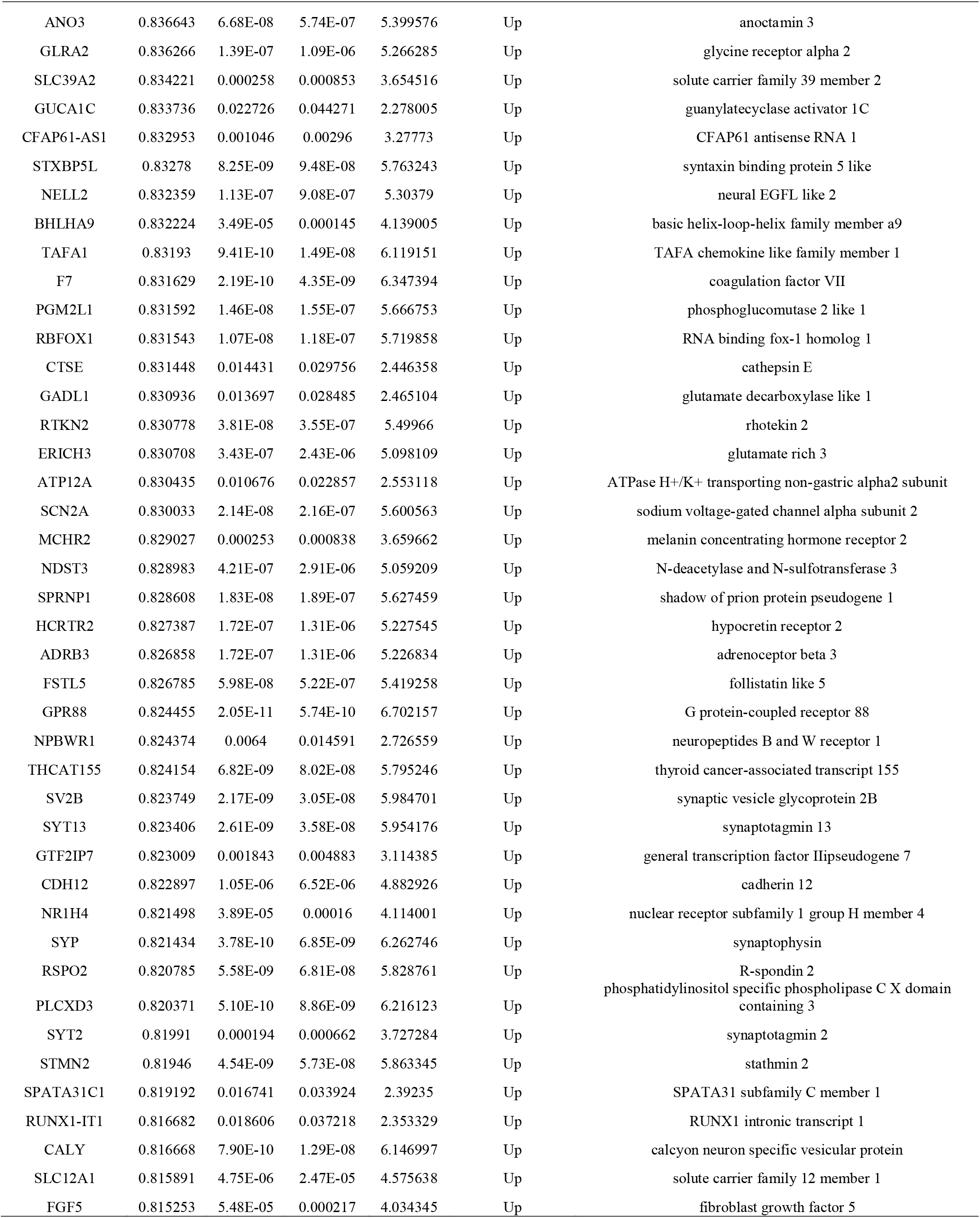
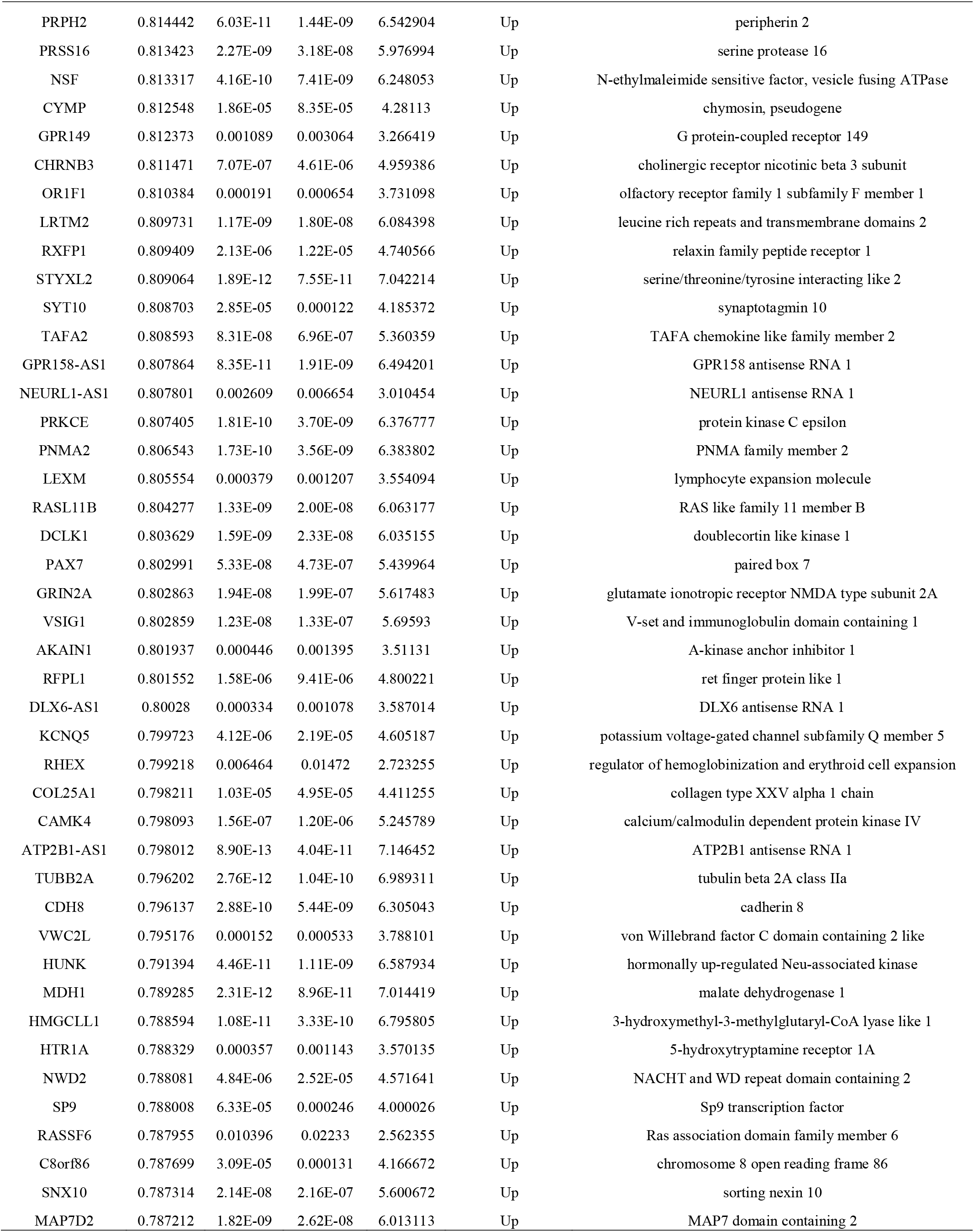
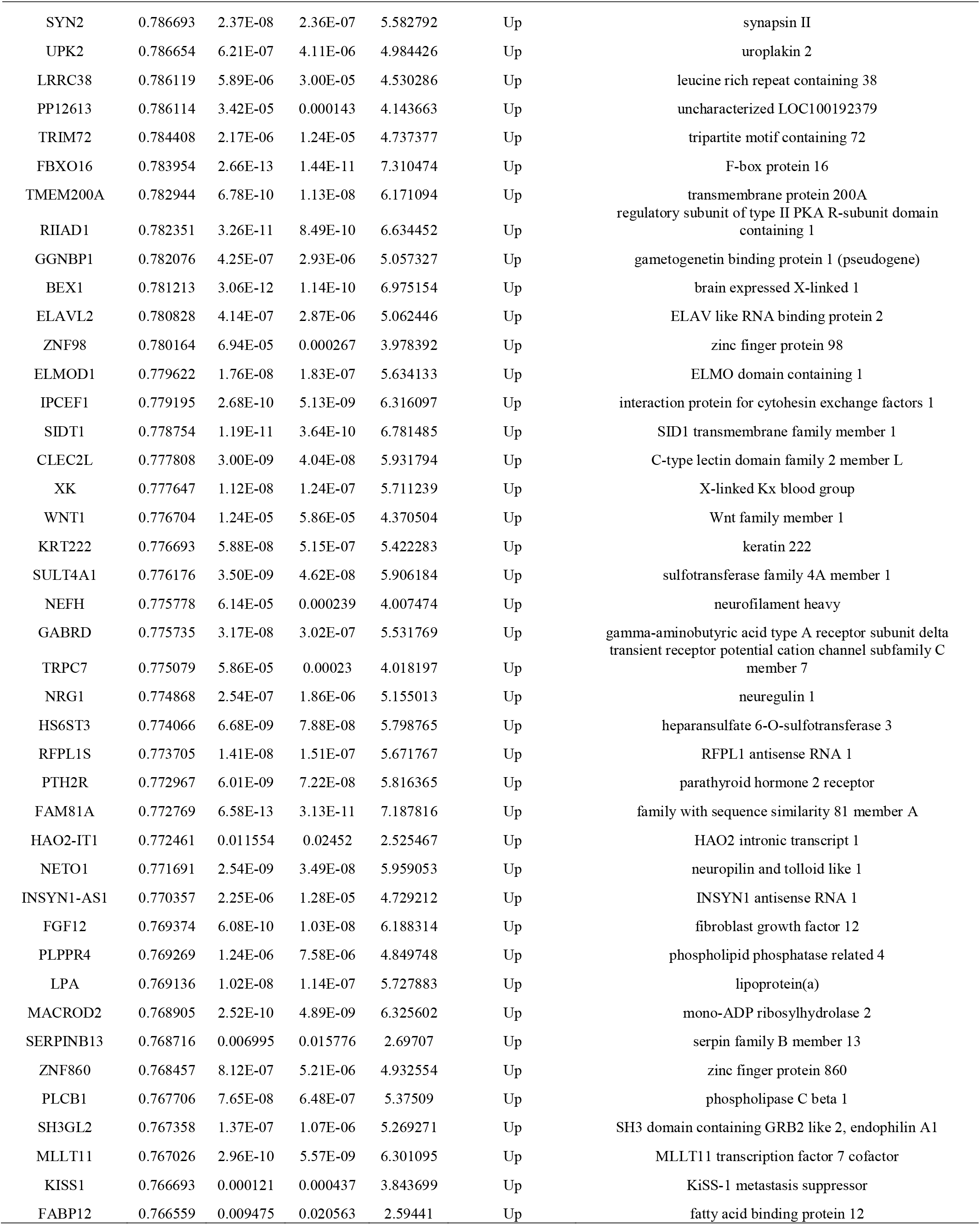
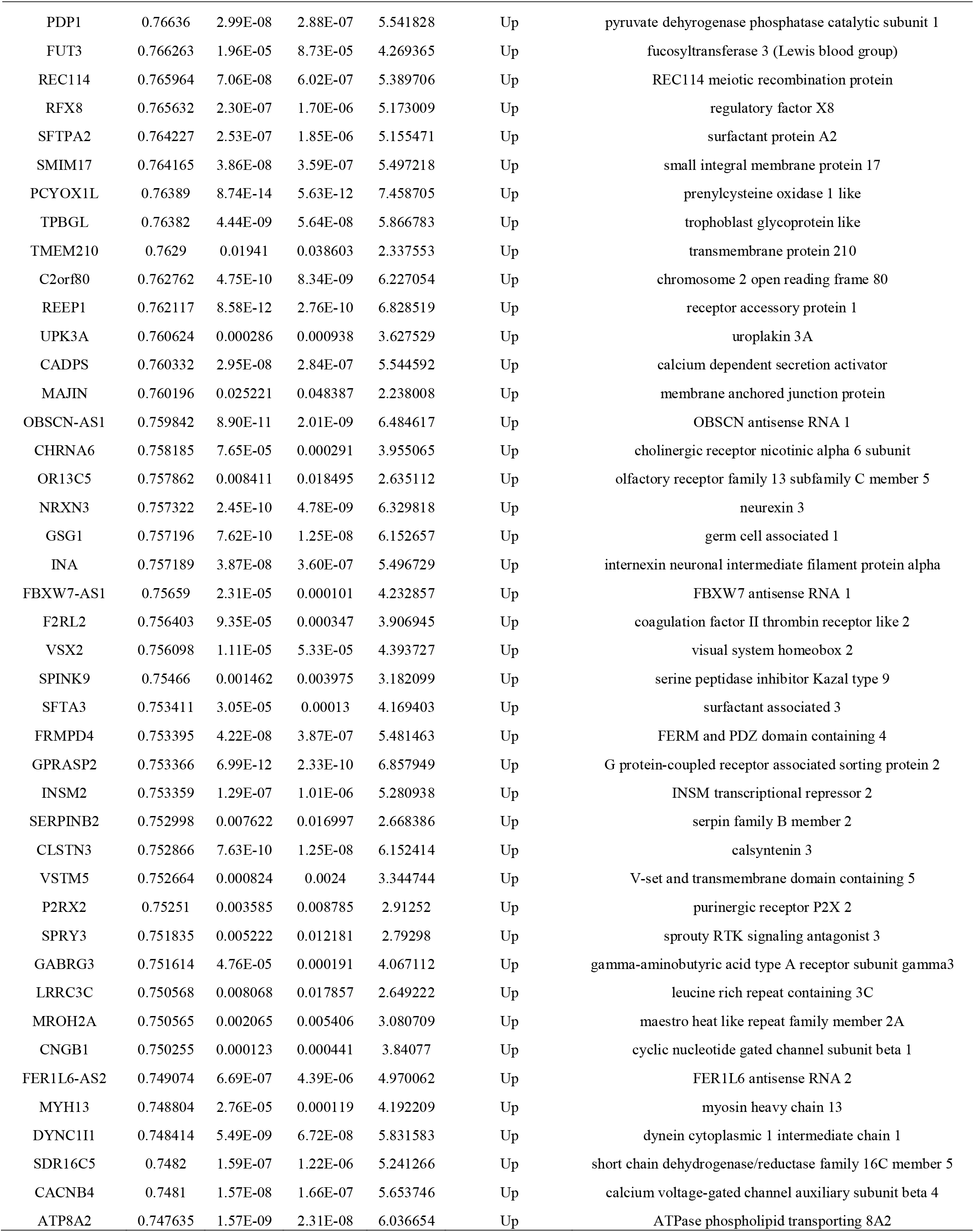
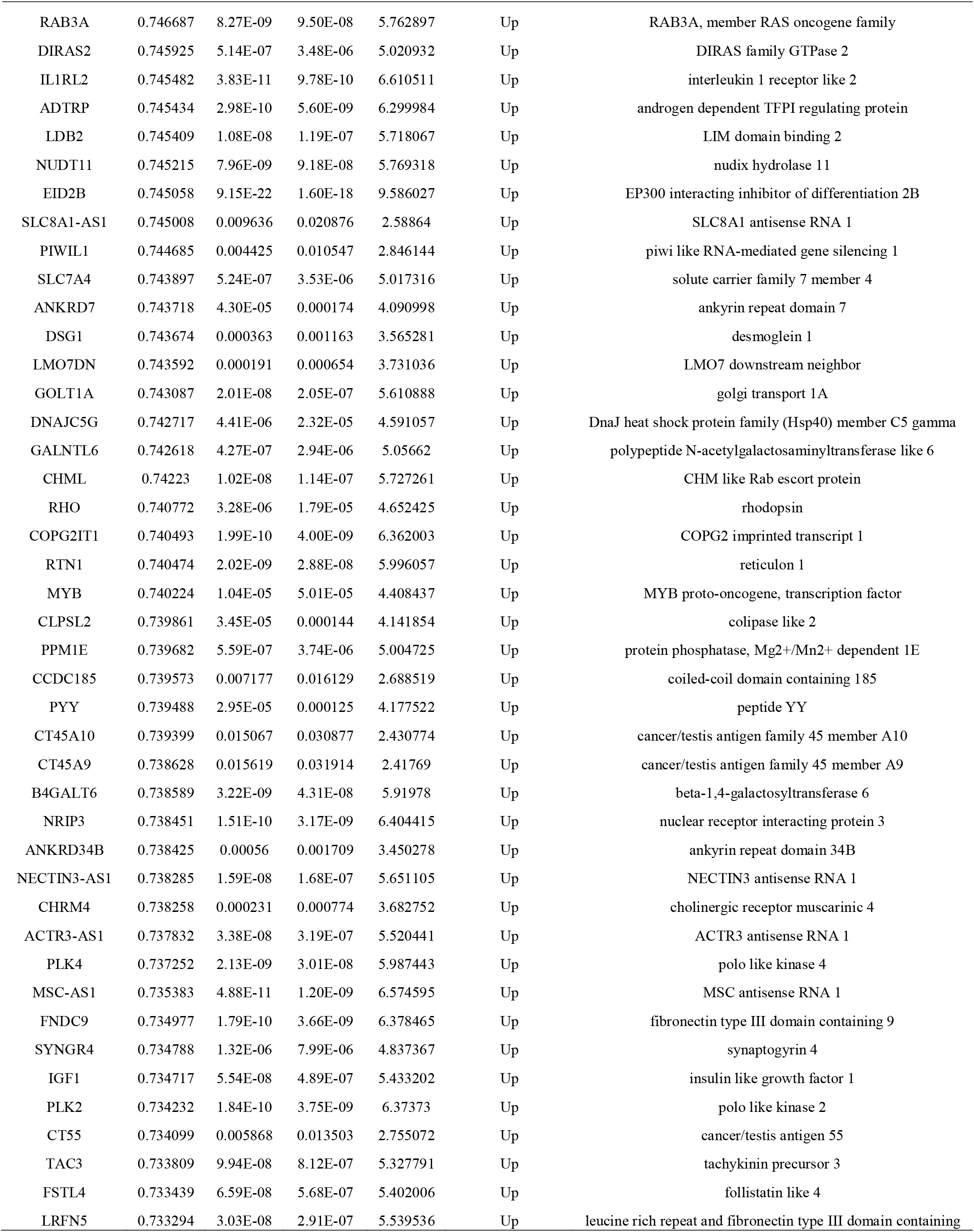
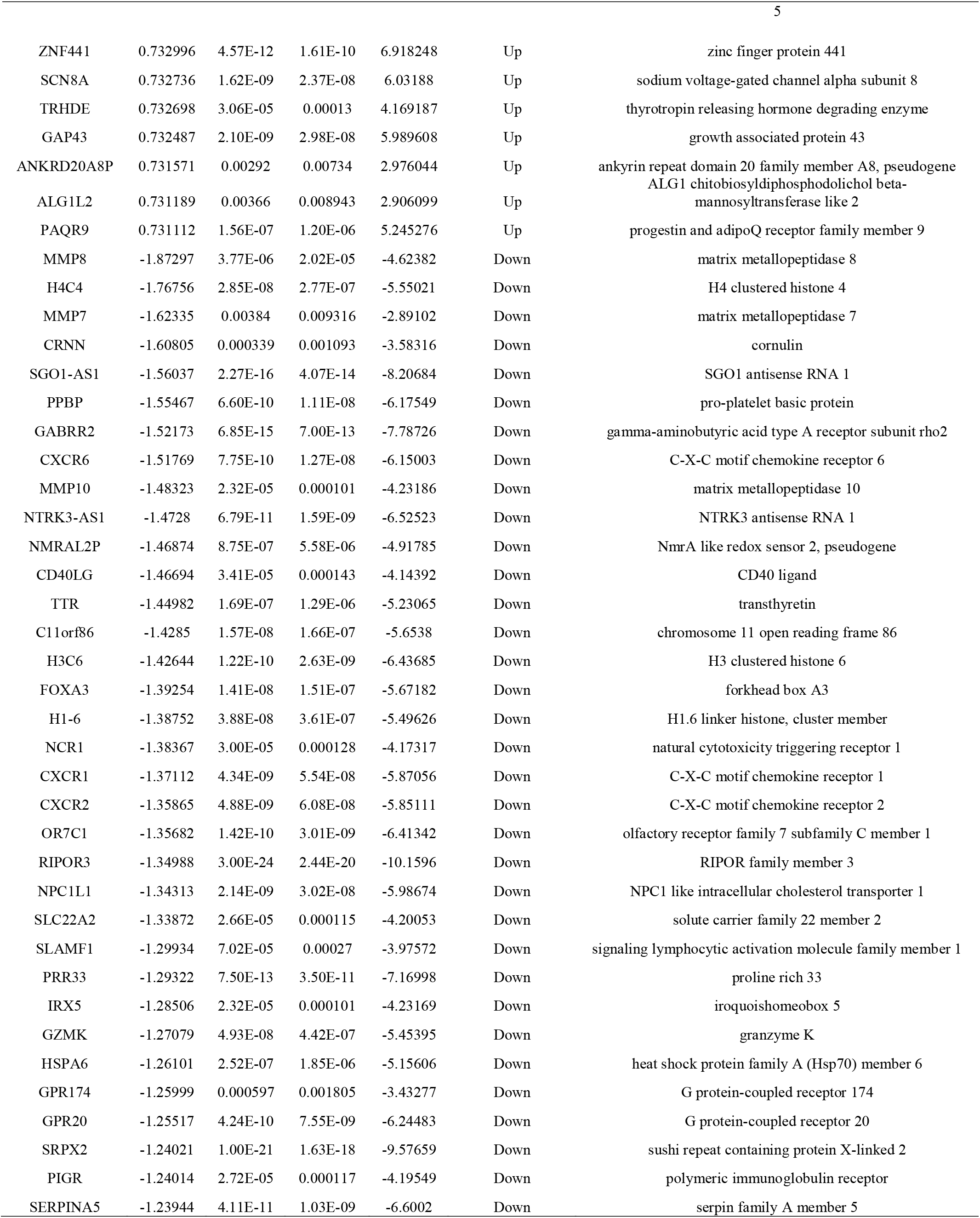
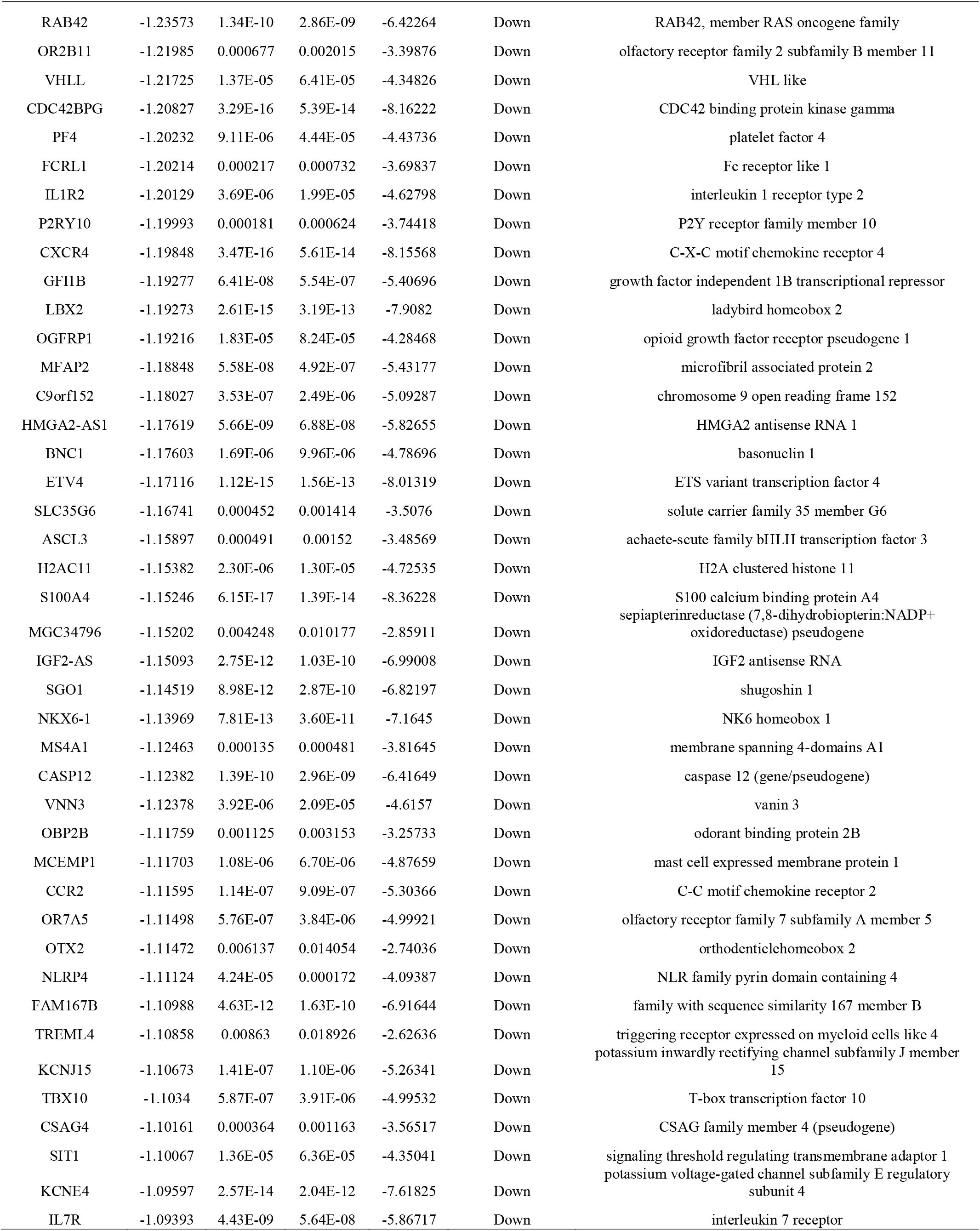
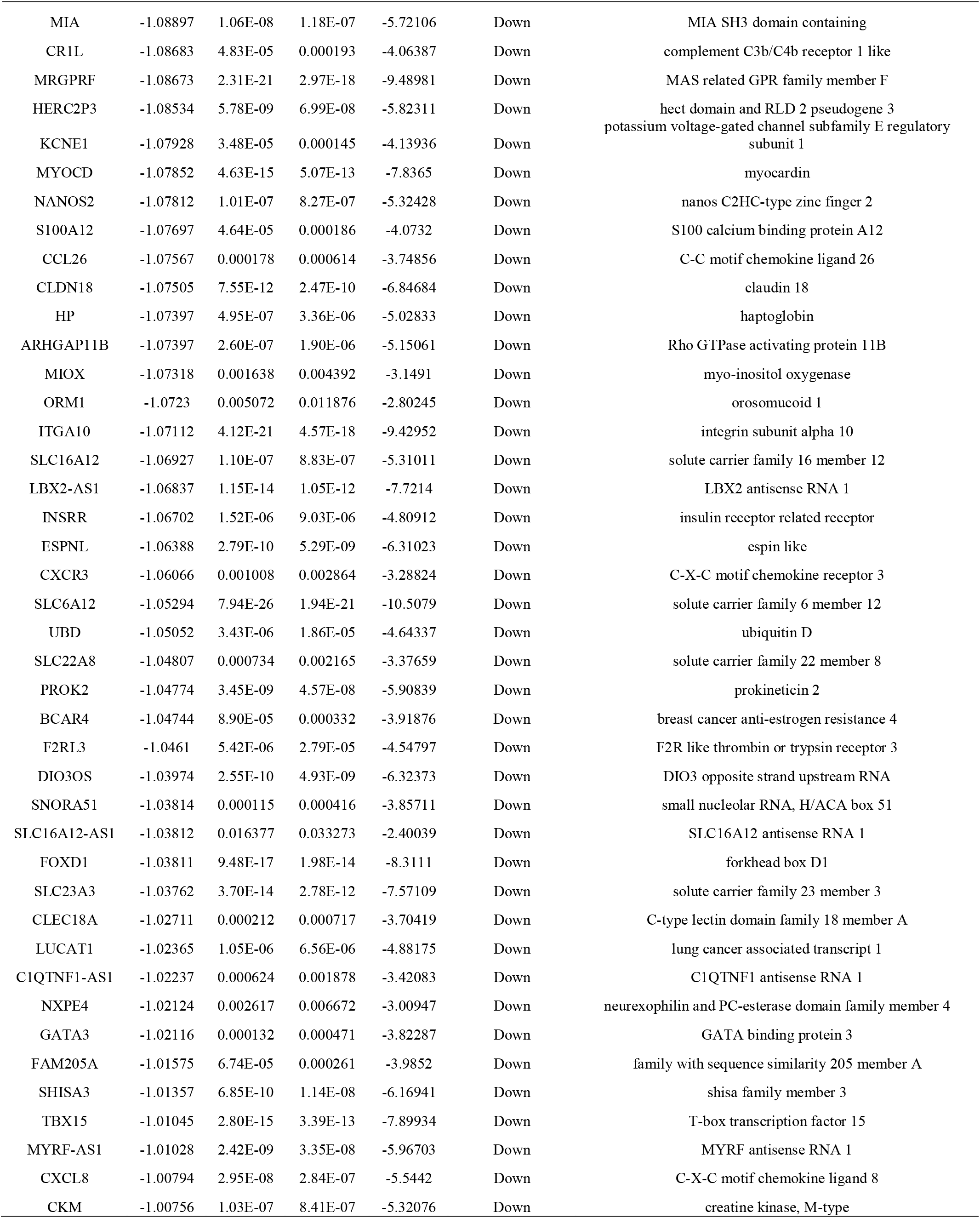
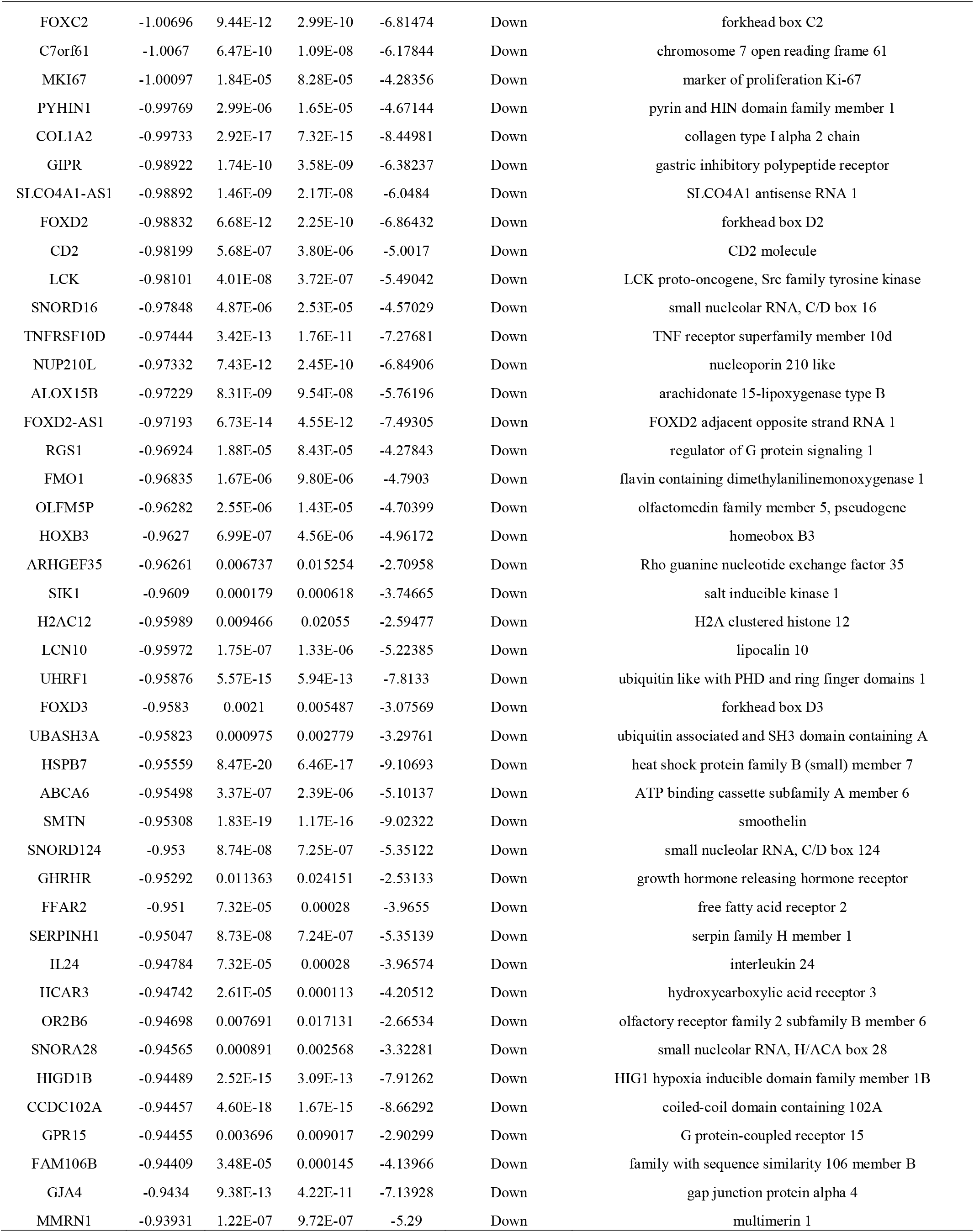
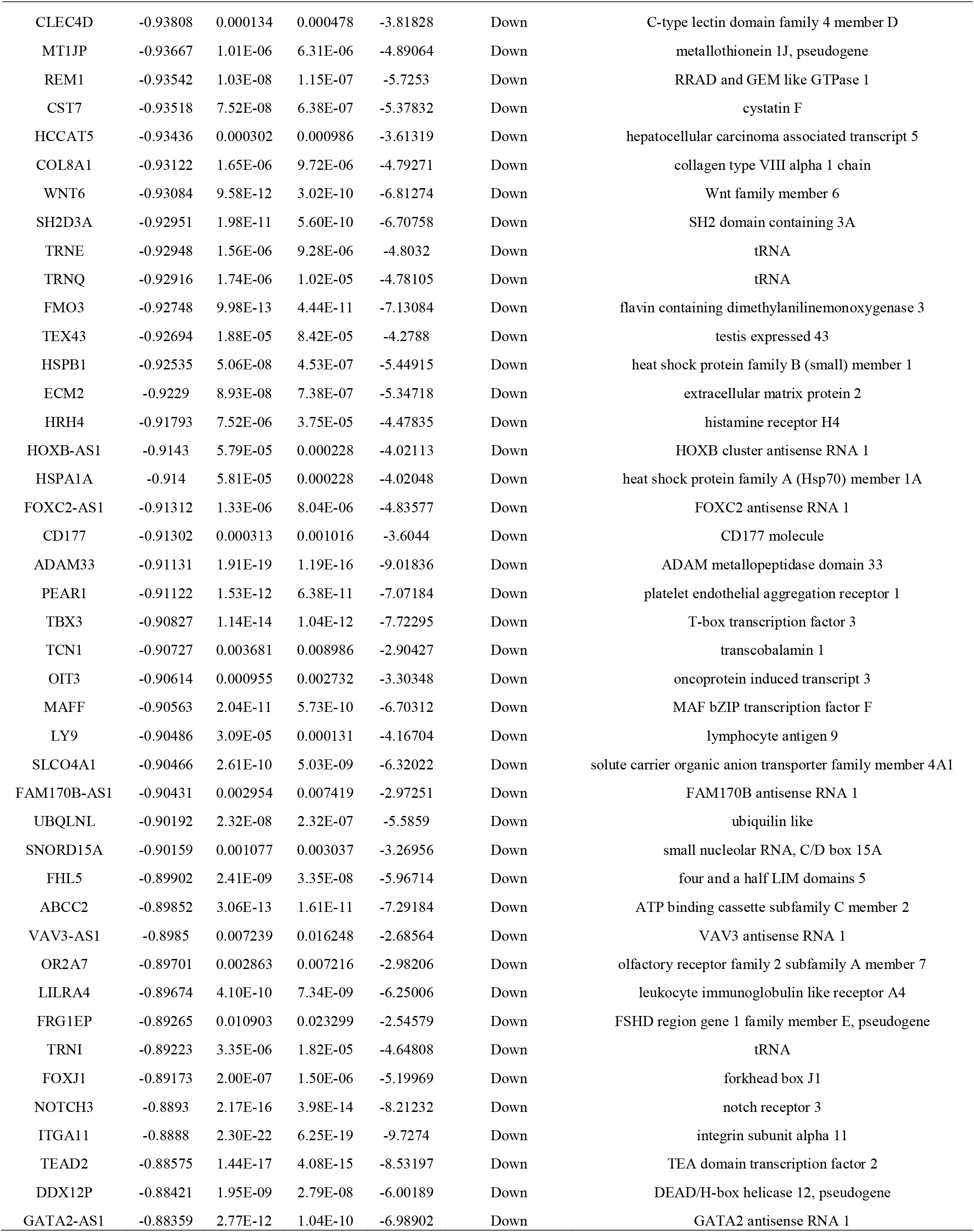
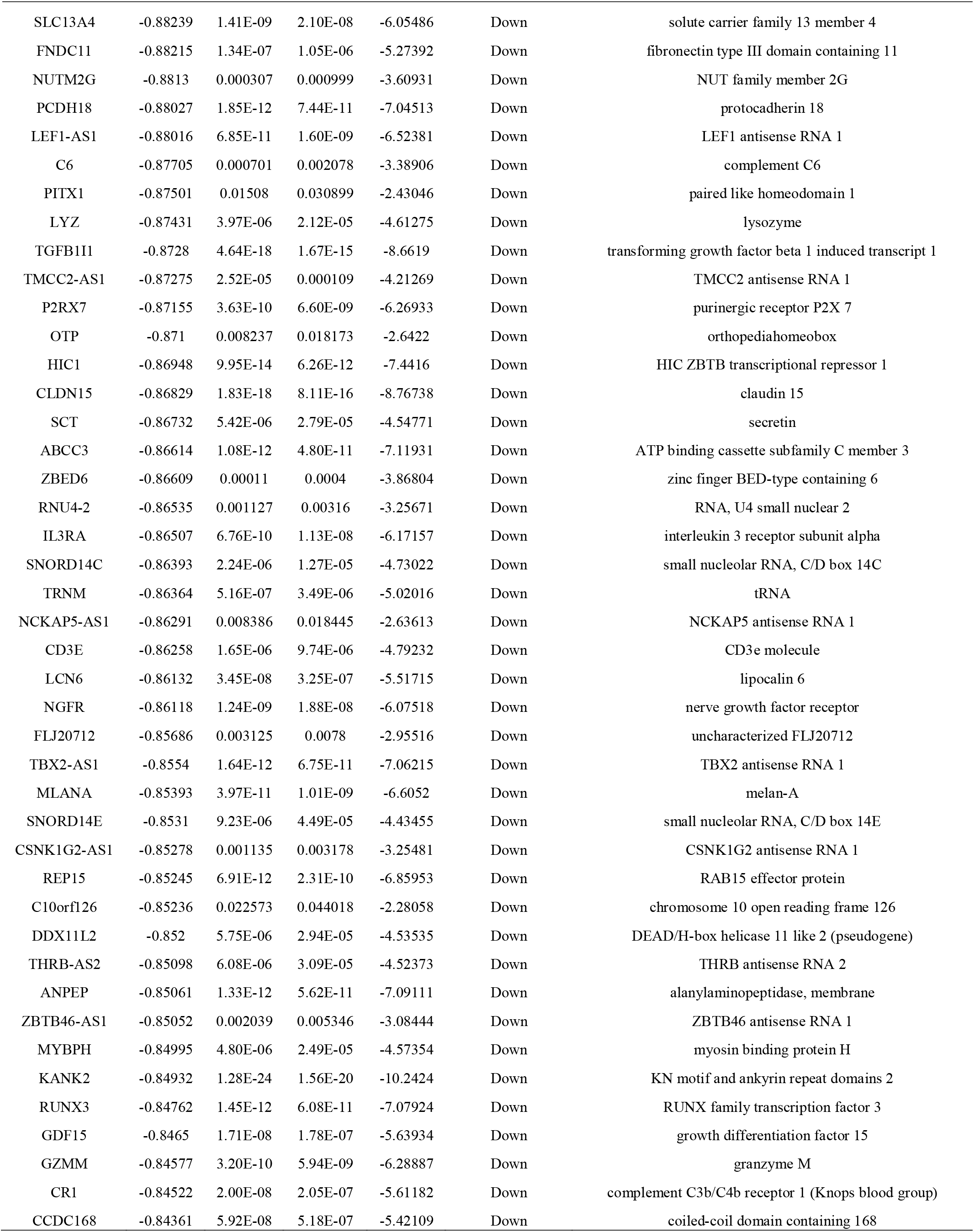
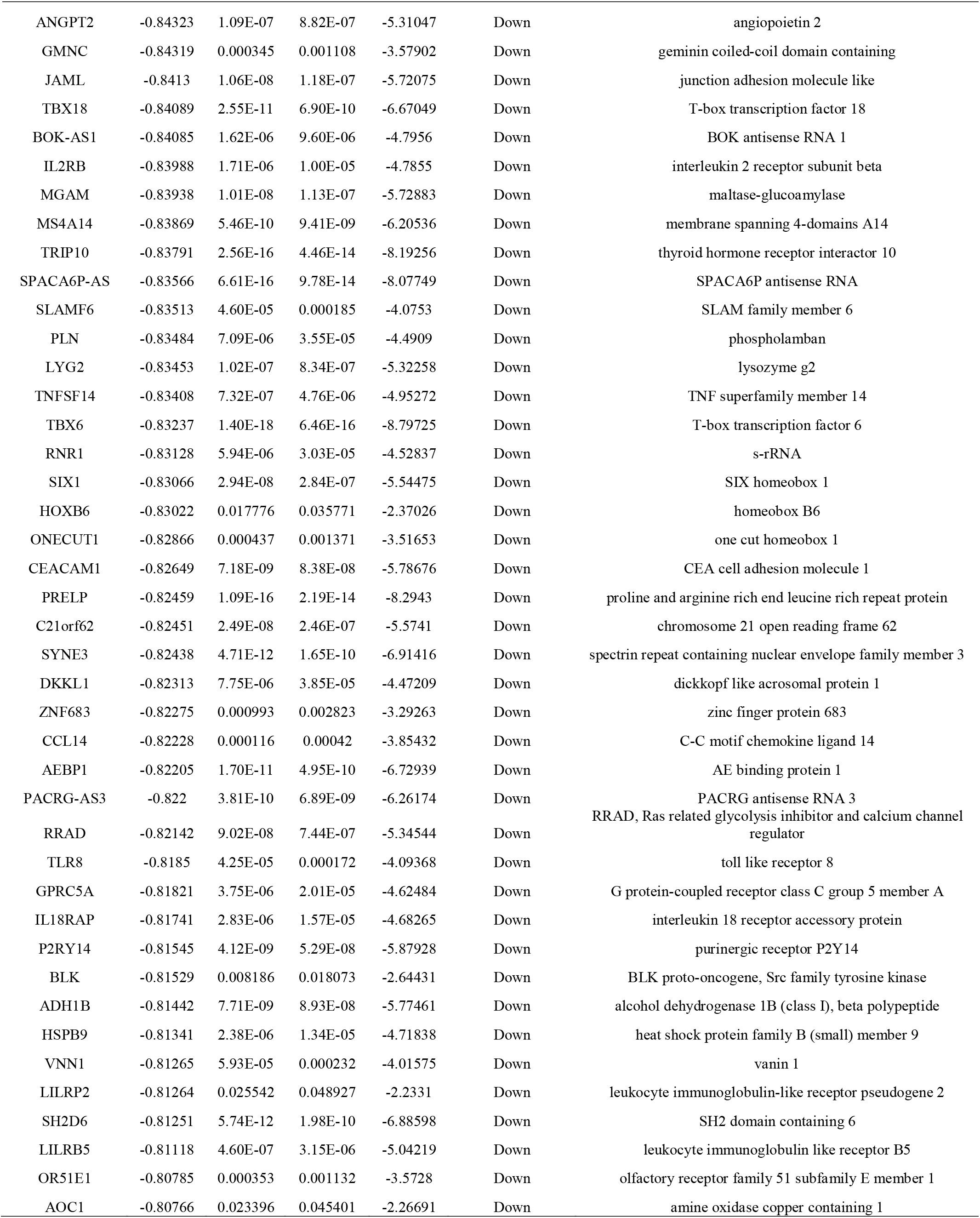
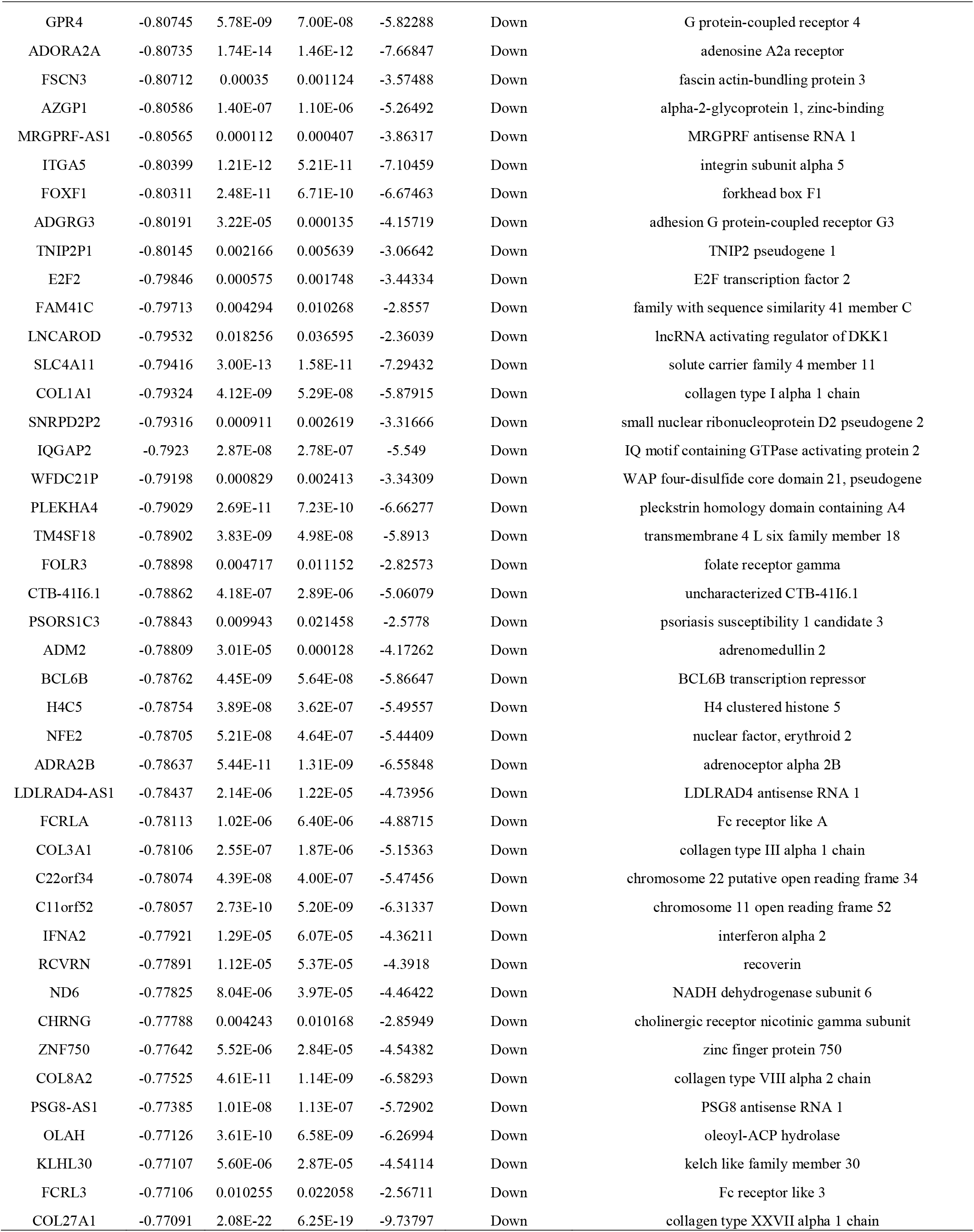
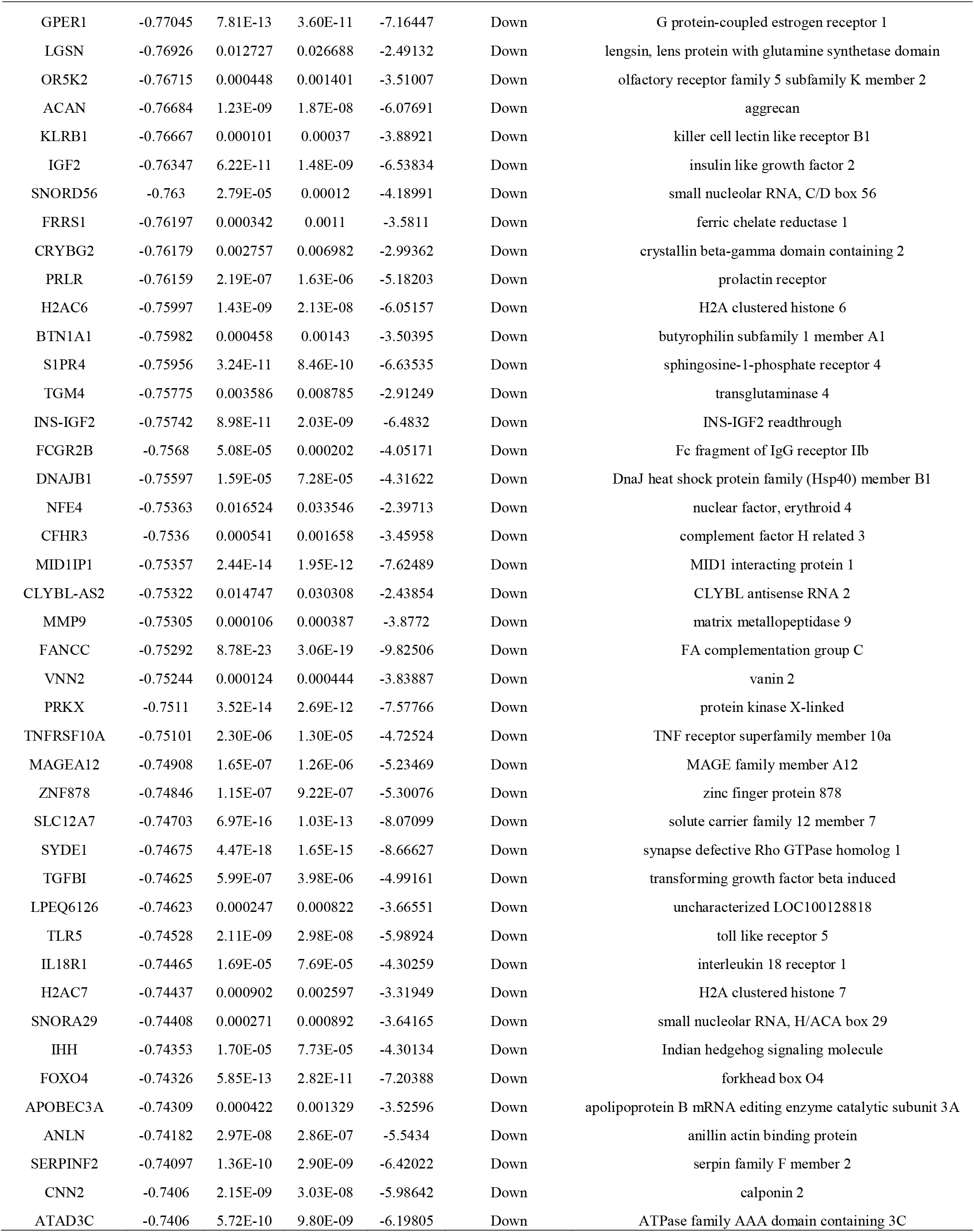
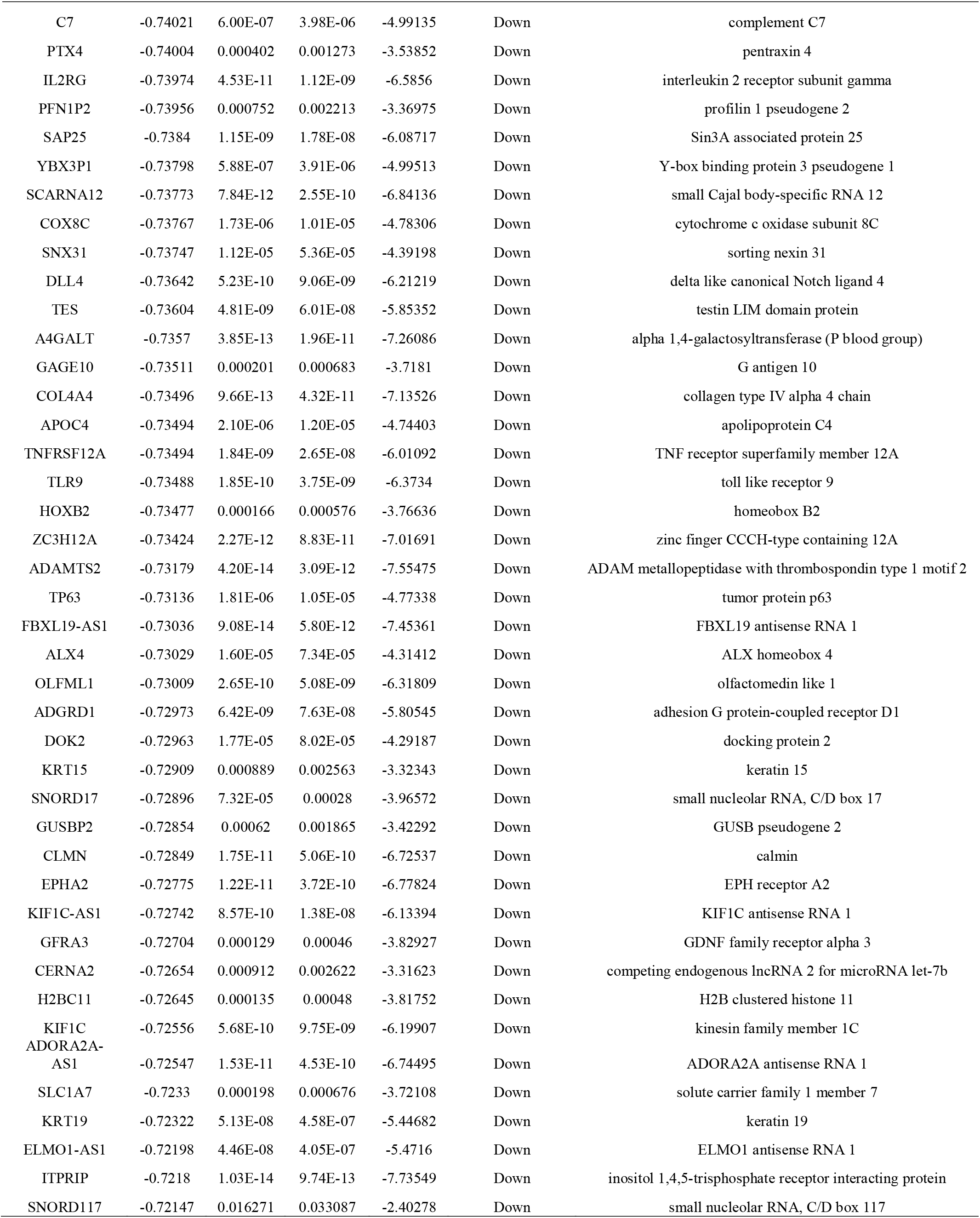
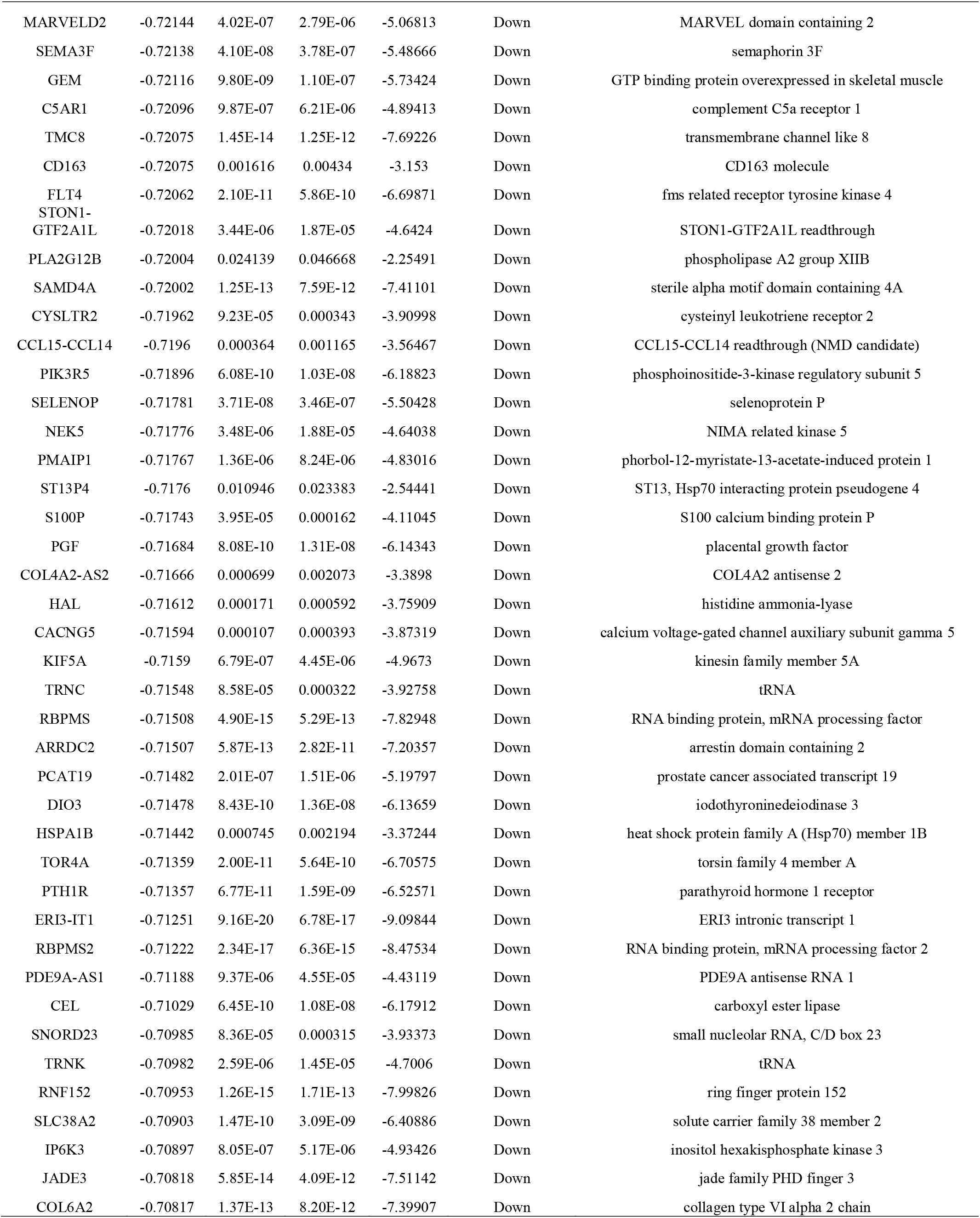
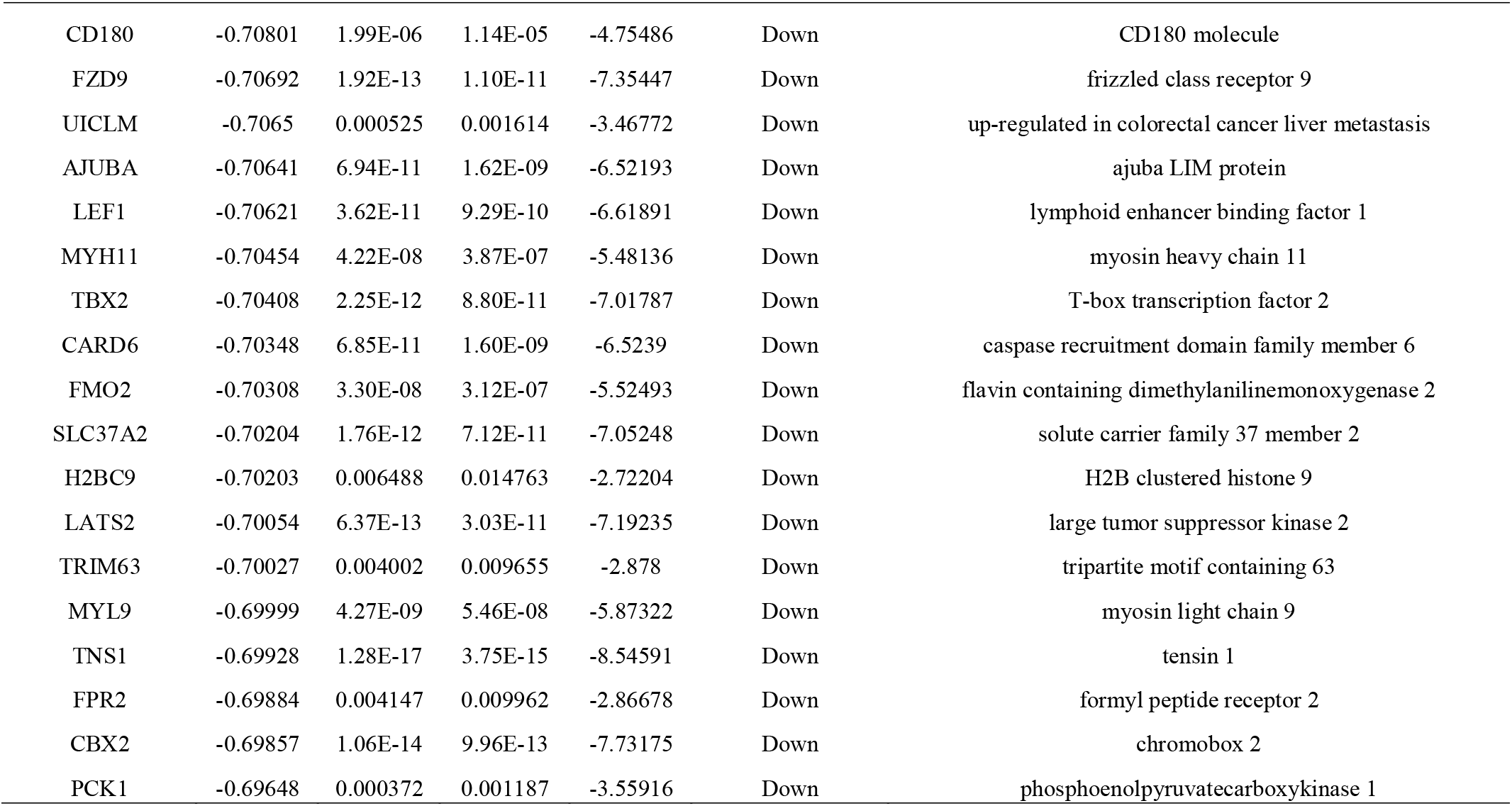
The statistical metrics for key differentially expressed genes (DEGs)

**Fig. 1.**
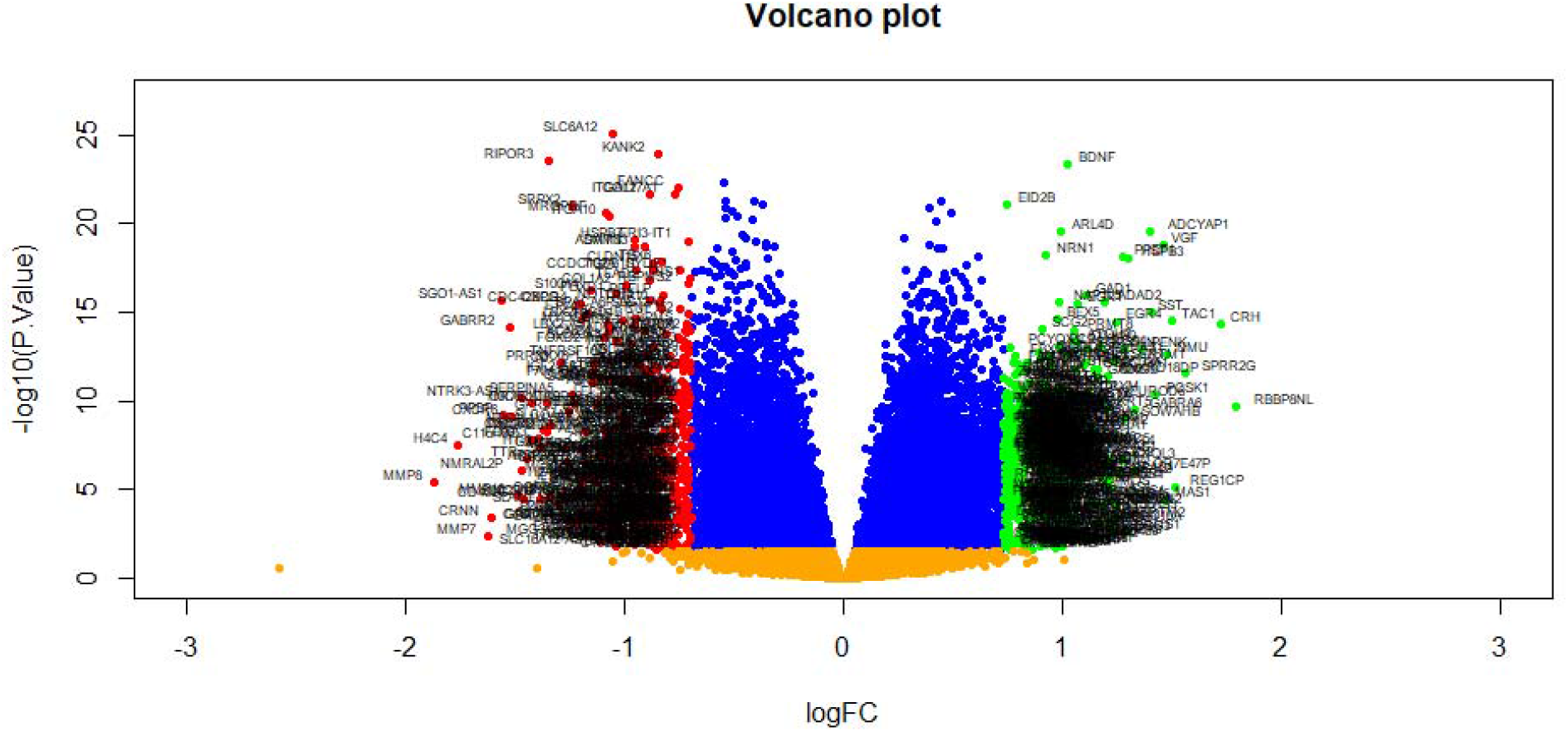
Volcano plot of differentially expressed genes. Genes with a significant change of more than two-fold were selected. Green dot represented up regulated significant genes and red dot represented down regulated significant genes.

**Fig. 2.**
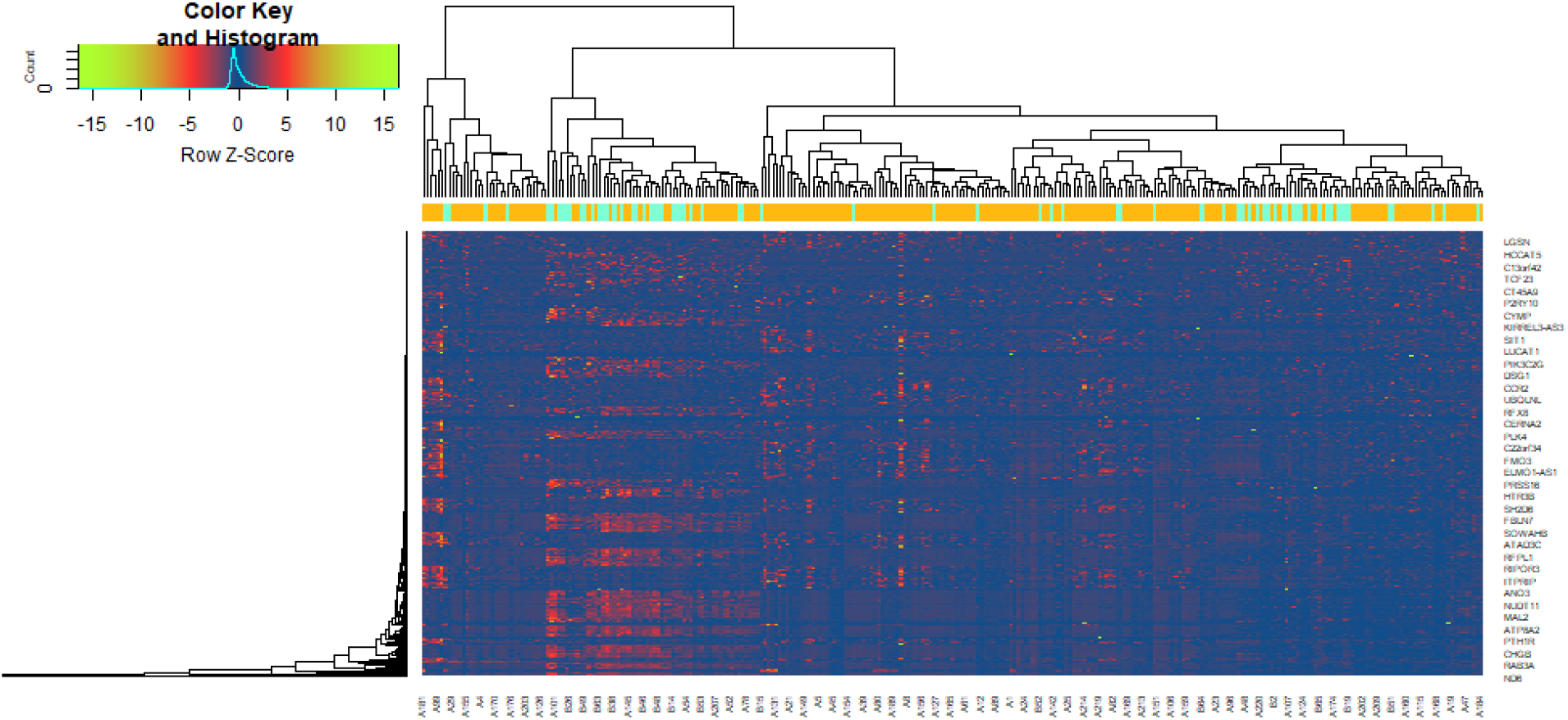
Heat map of differentially expressed genes. Legend on the top left indicate log fold change of genes. (A1 – A219= AD; B1 – B70 = healthy controls)

### GO and REACTOME pathway enrichment analysis of DEGs

We performed GO and REACTOME pathway enrichment analysis to investigate the functions of DEGs using g:Profiler. The top GO (Table 2) and REACTOME pathway (Table 3) terms for DEGs were shown. For BP, DEGs were mainly enriched in regulation of biological quality, multicellular organismal process, cell communication and response to stimulus. The CC analysis indicated that proteins encoded by DEGs were mostly located in the cell junction, cell periphery, plasma membrane and extracellular region. DEGs in molecular function (MF) were significantly associated with channel activity, transmembranesignaling receptor activity, signaling receptor activity and G protein-coupled receptor activity. REACTOME pathway enrichment analysis showed that neuronal system, neurotransmitter receptors and postsynaptic signal transmission, GPCR ligand binding and signal transduction were significantly enriched in DEGs.

**Table 2.**
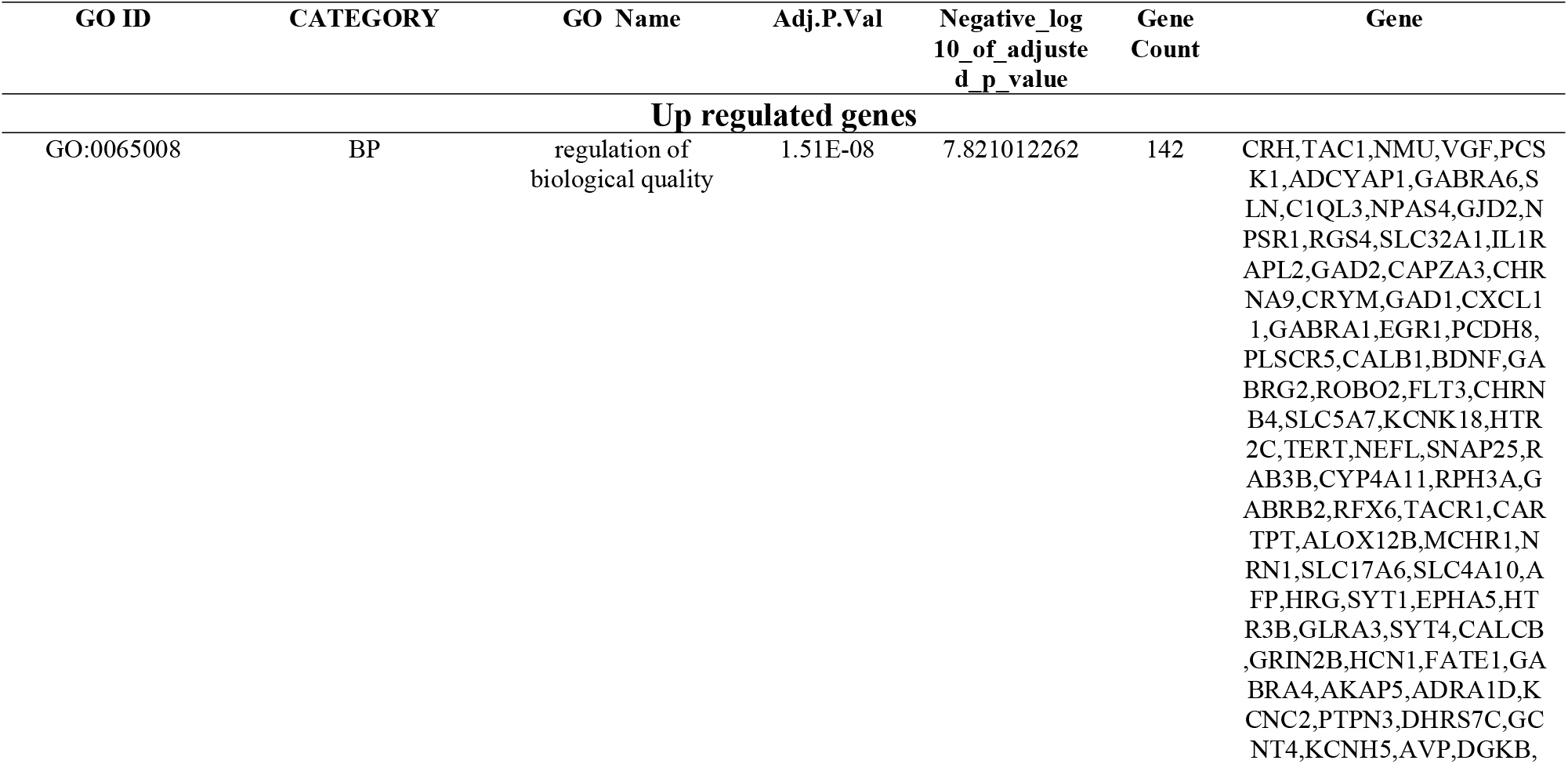
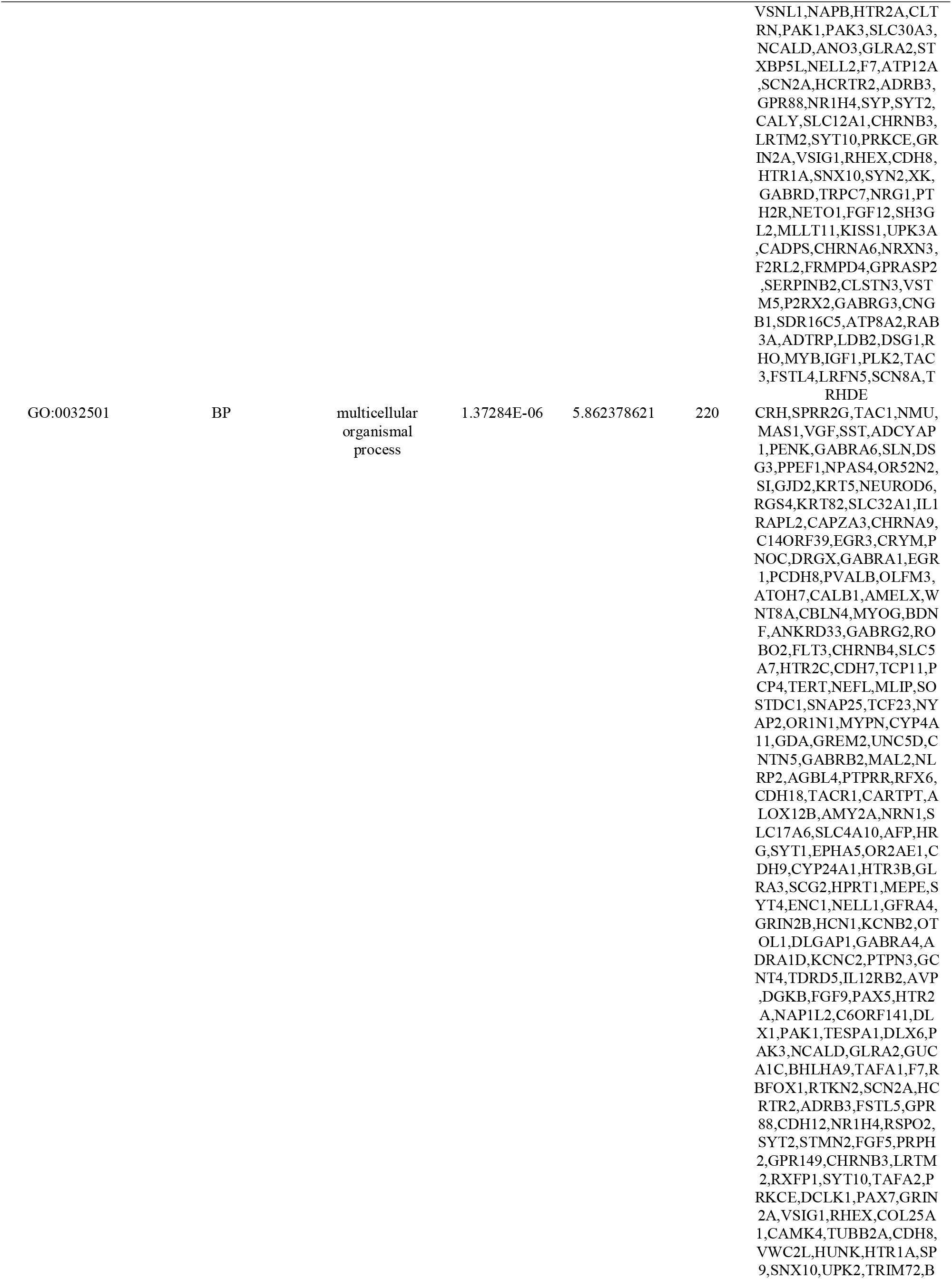
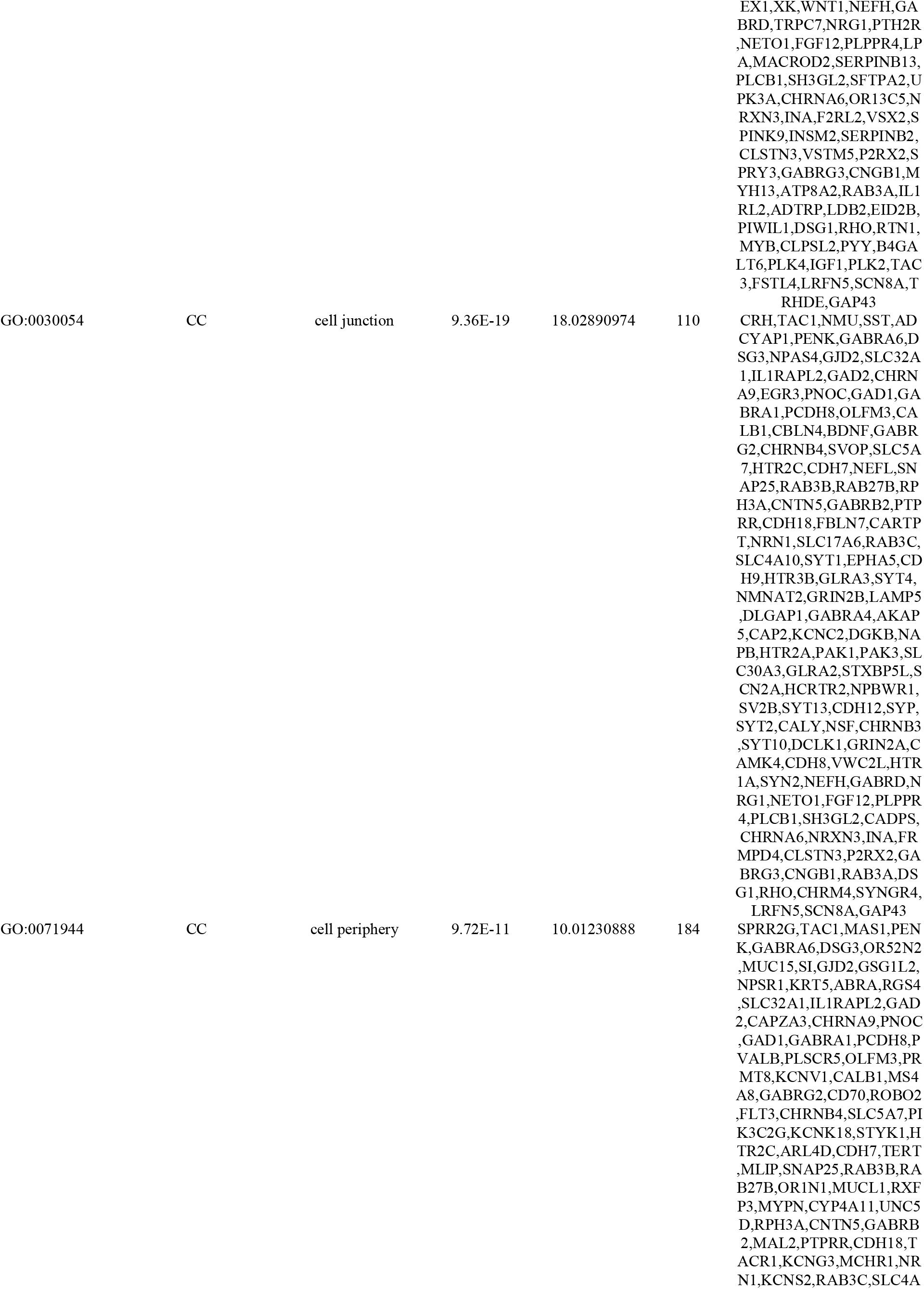
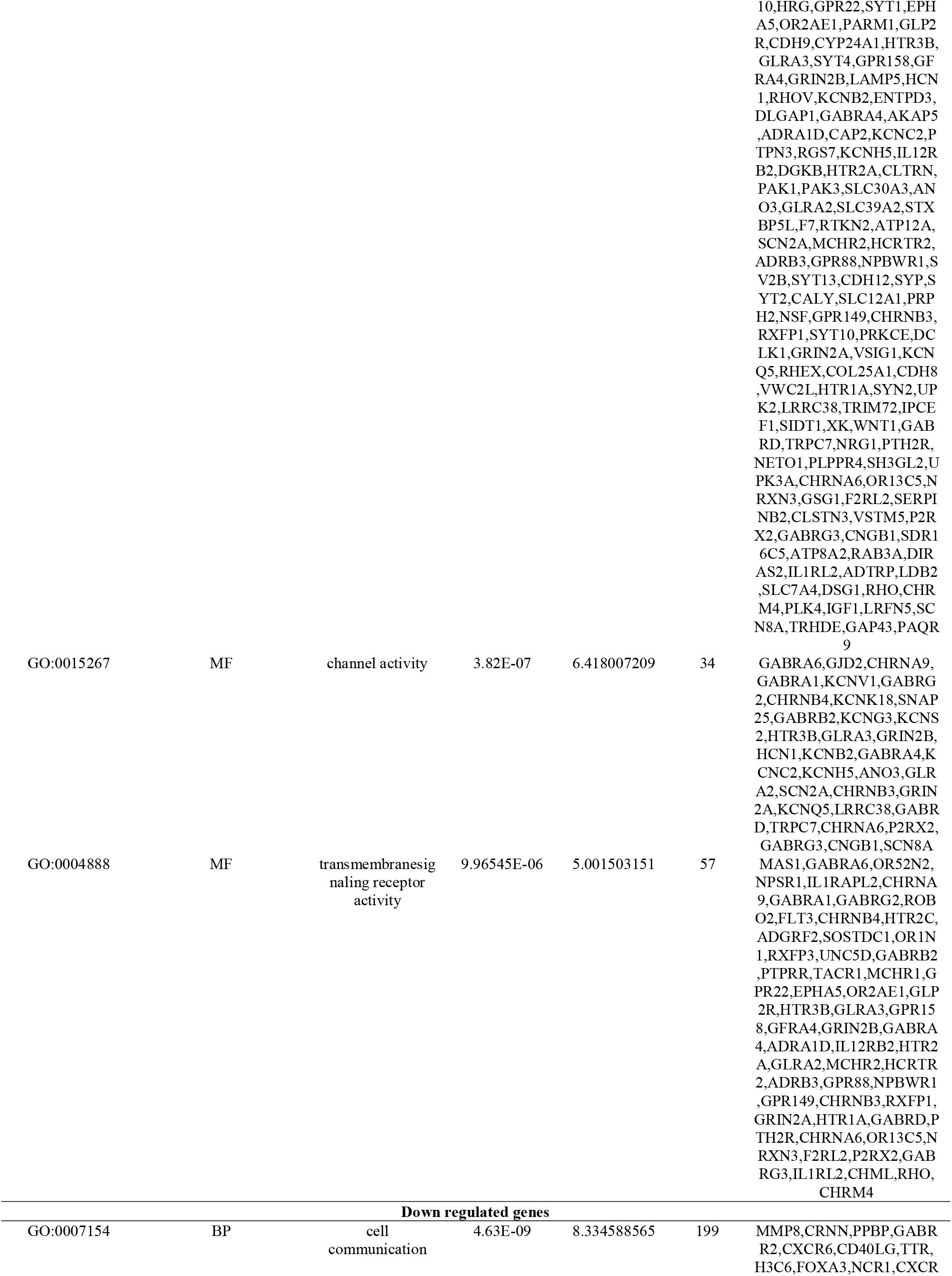
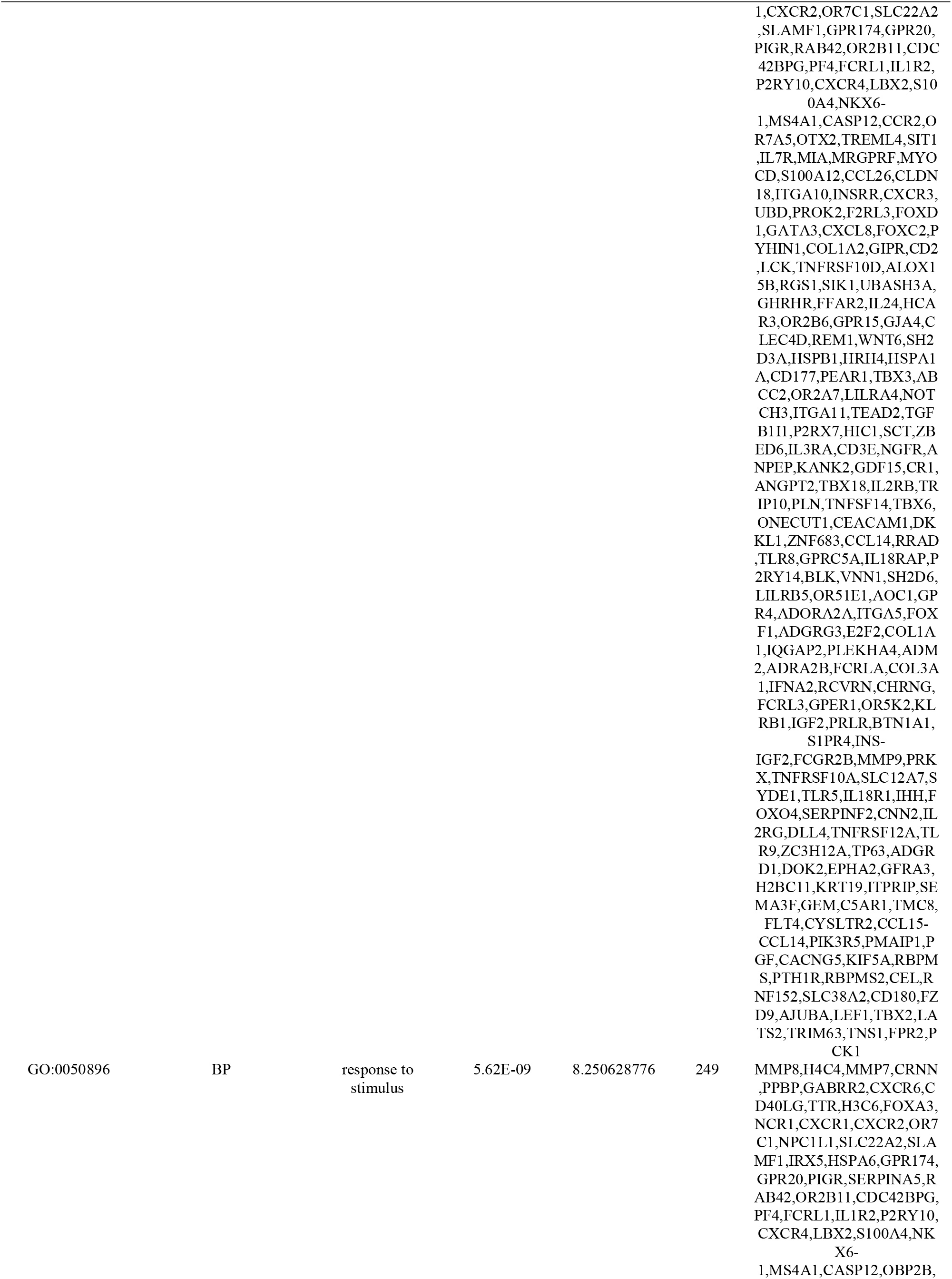
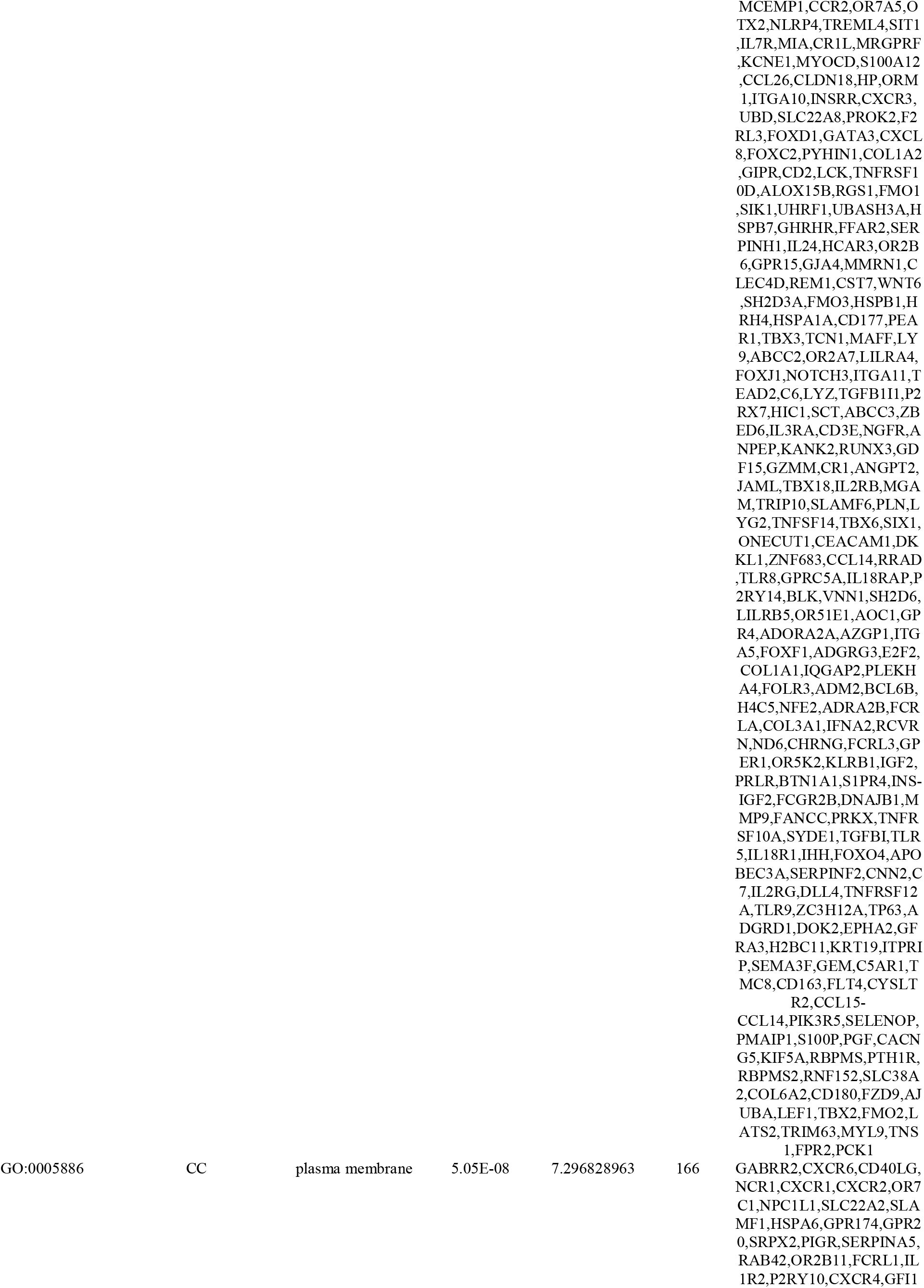
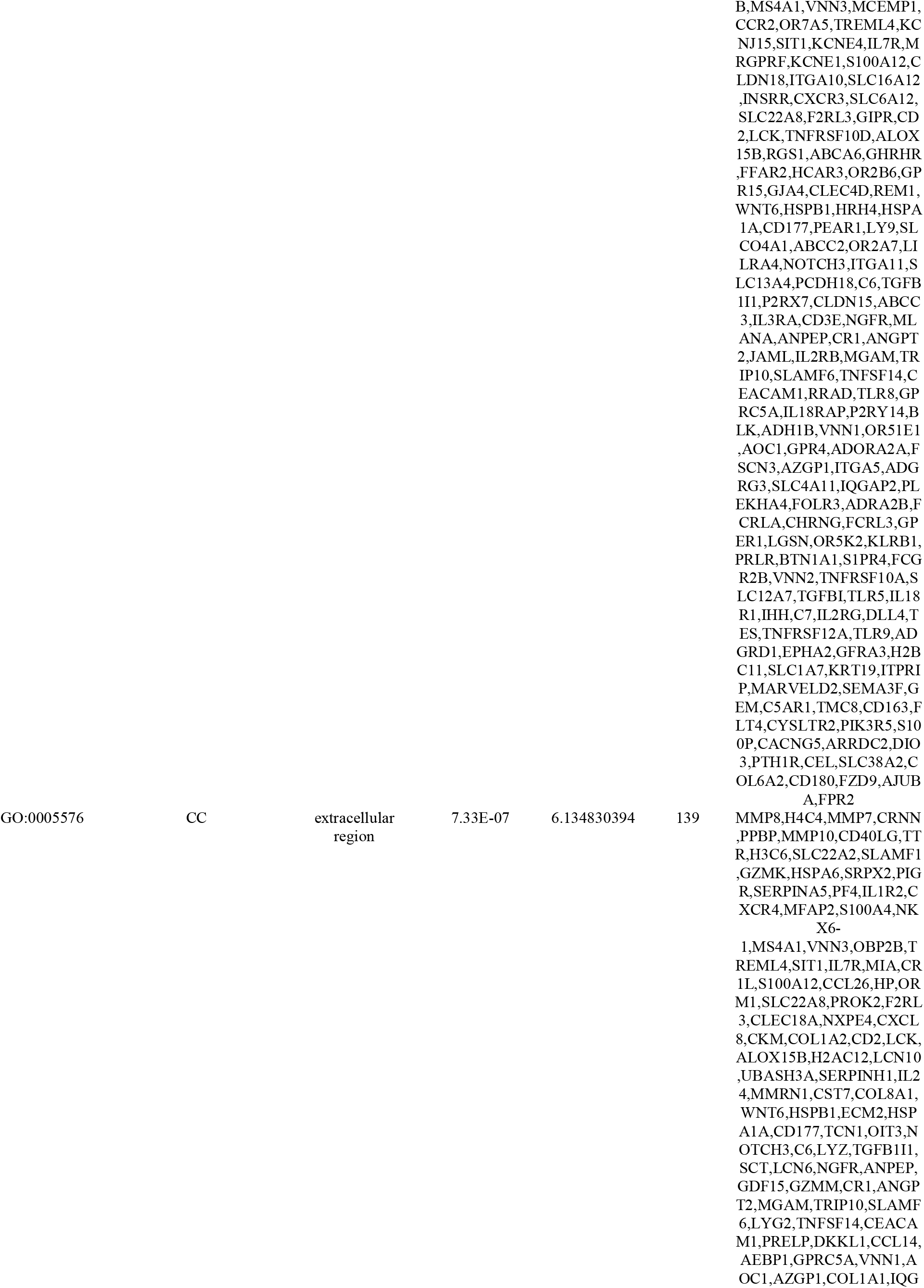
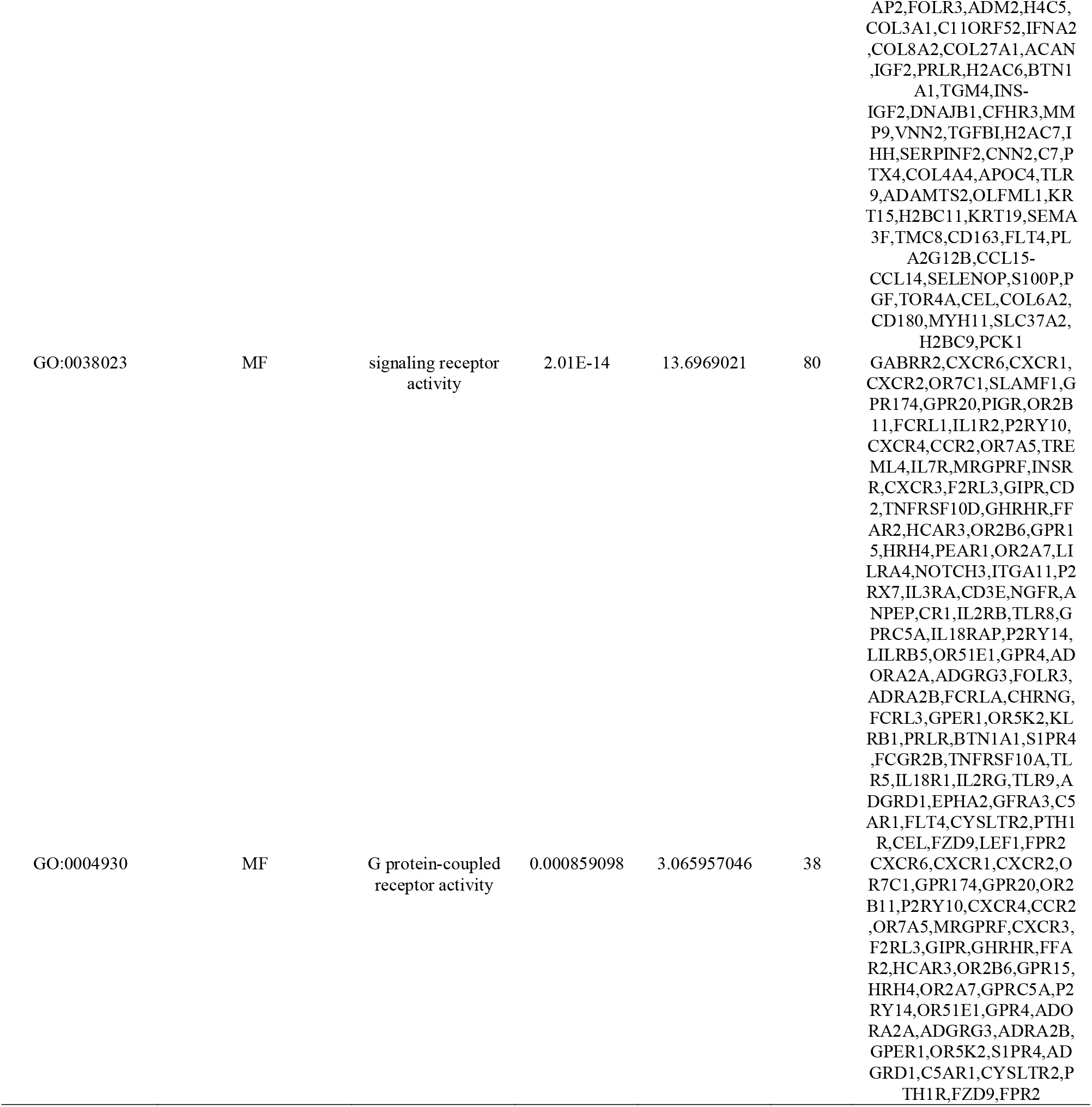
The enriched GO terms of the up and down regulated differentially expressed genes

**Table 3.**
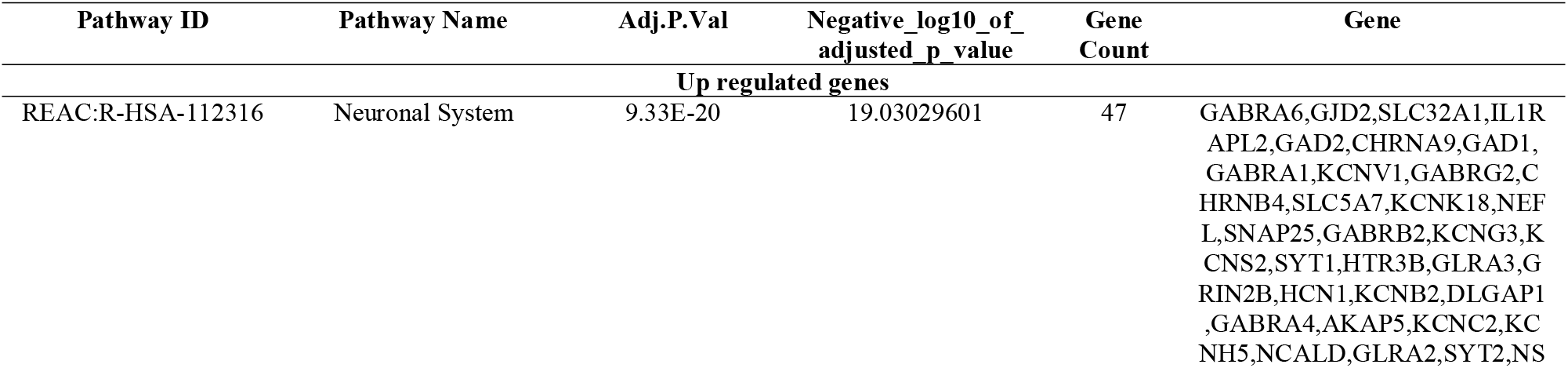
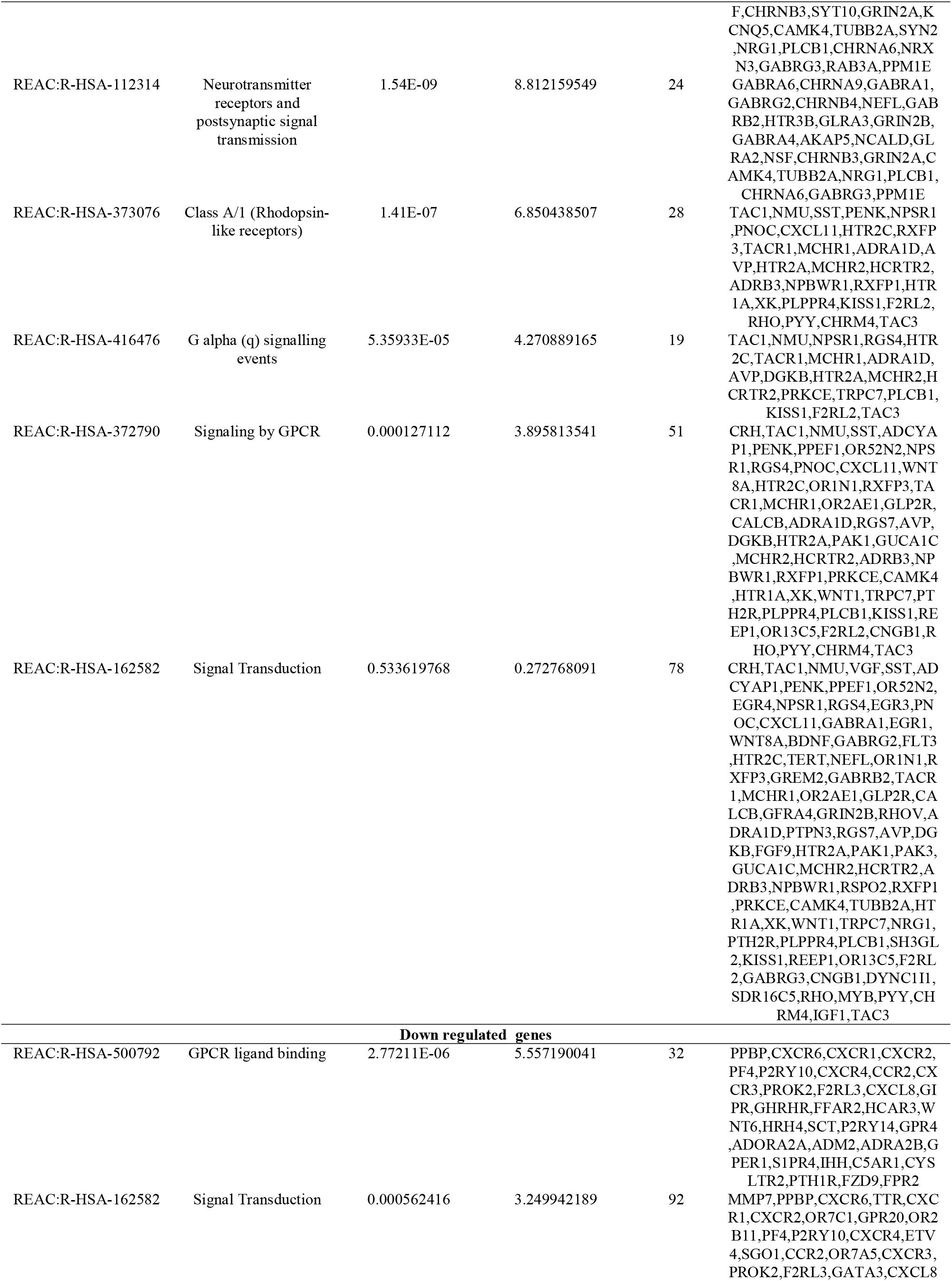
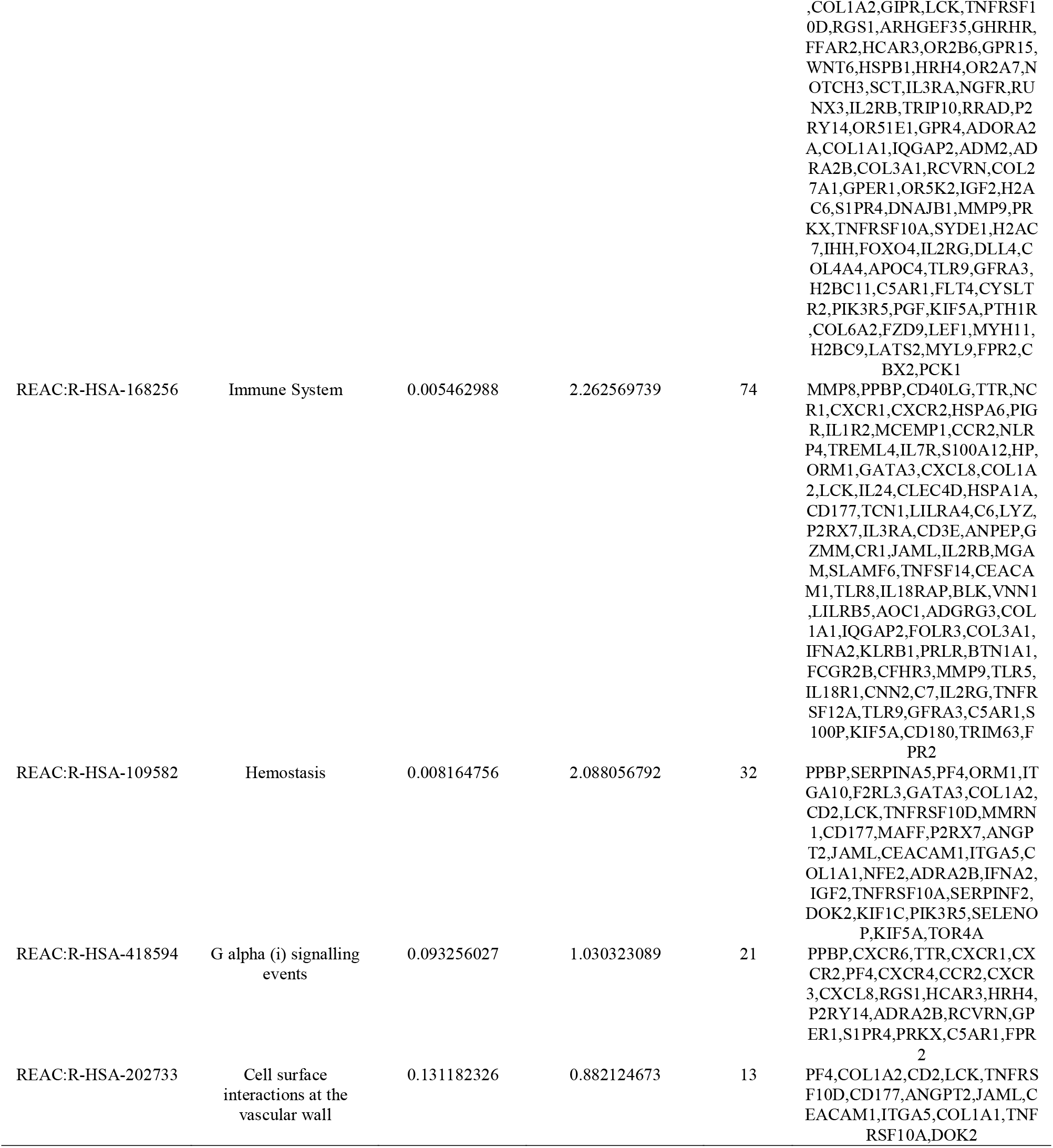
The enriched pathway terms of the up and down regulated differentially expressed genes

### PPI network establishment and modules selection

The PPI network of DEGs consisted of 6612 nodes and 13538 edges constructed in the HIPPIE interactome database and visualized using Cytoscape software (Fig.3). Based on the HIPPIE interactome database, the DEGs with the highest PPI scores identified by the four centrality methods are shown in Table 4. After repeated genes were removed, the hub genes (shown in Fig. 3, highlighted in green color for up regulated genes and red color for down regulated genes and shaped in round) were obtained using the four centrality methods, including PAK1, ELAVL2, NSF, HTR2C, TERT, UBD, MKI67, HSPB1, PYHIN1 and TES. A two significant modules were constructed from the PPI network of the DEGs using PEWCC1, including module 1 had 16 nodes and 38 edges (Fig. 4A) and module 2 had 47 nodes and 96 edges (Fig. 4B). GO and pathway enrichment analysis showed that genes in this module were markedly enriched in multicellular organismal process, neuronal system, regulation of biological quality, signal transduction, cell communication, plasma membrane, response to stimulus and cell communication.

**Fig. 3.**
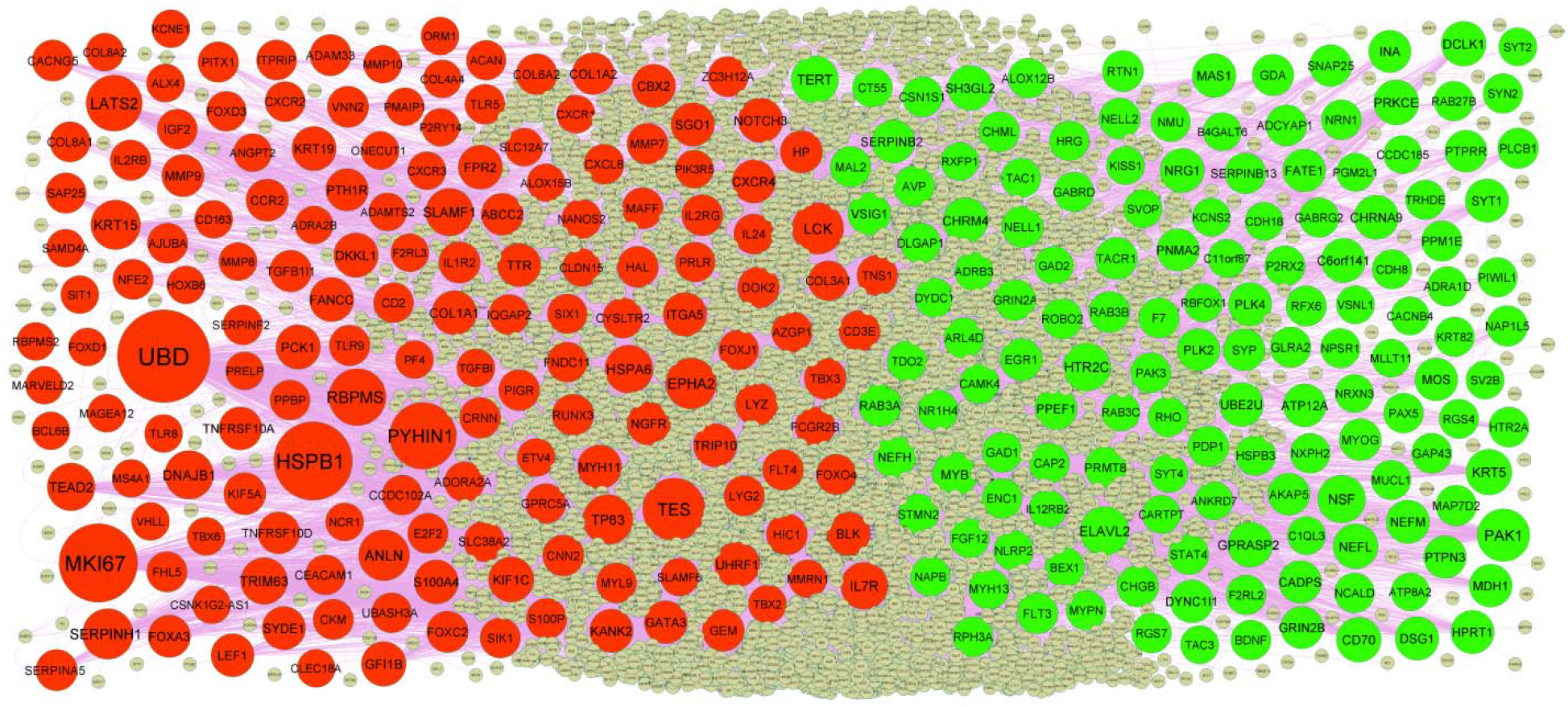
PPI network of DEGs. The PPI network of DEGs was constructed using Cytoscap. Up regulated genes are marked in green; down regulated genes are marked in red

**Table 4.**
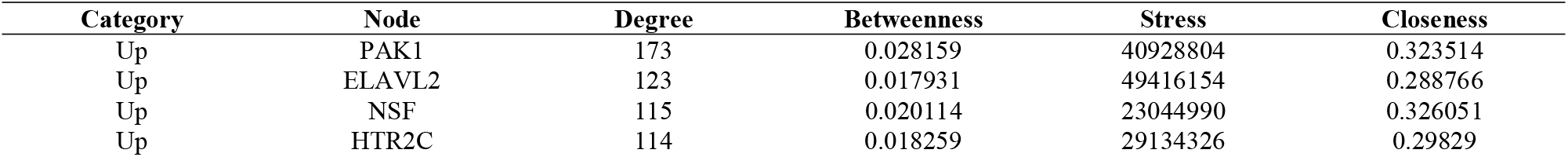
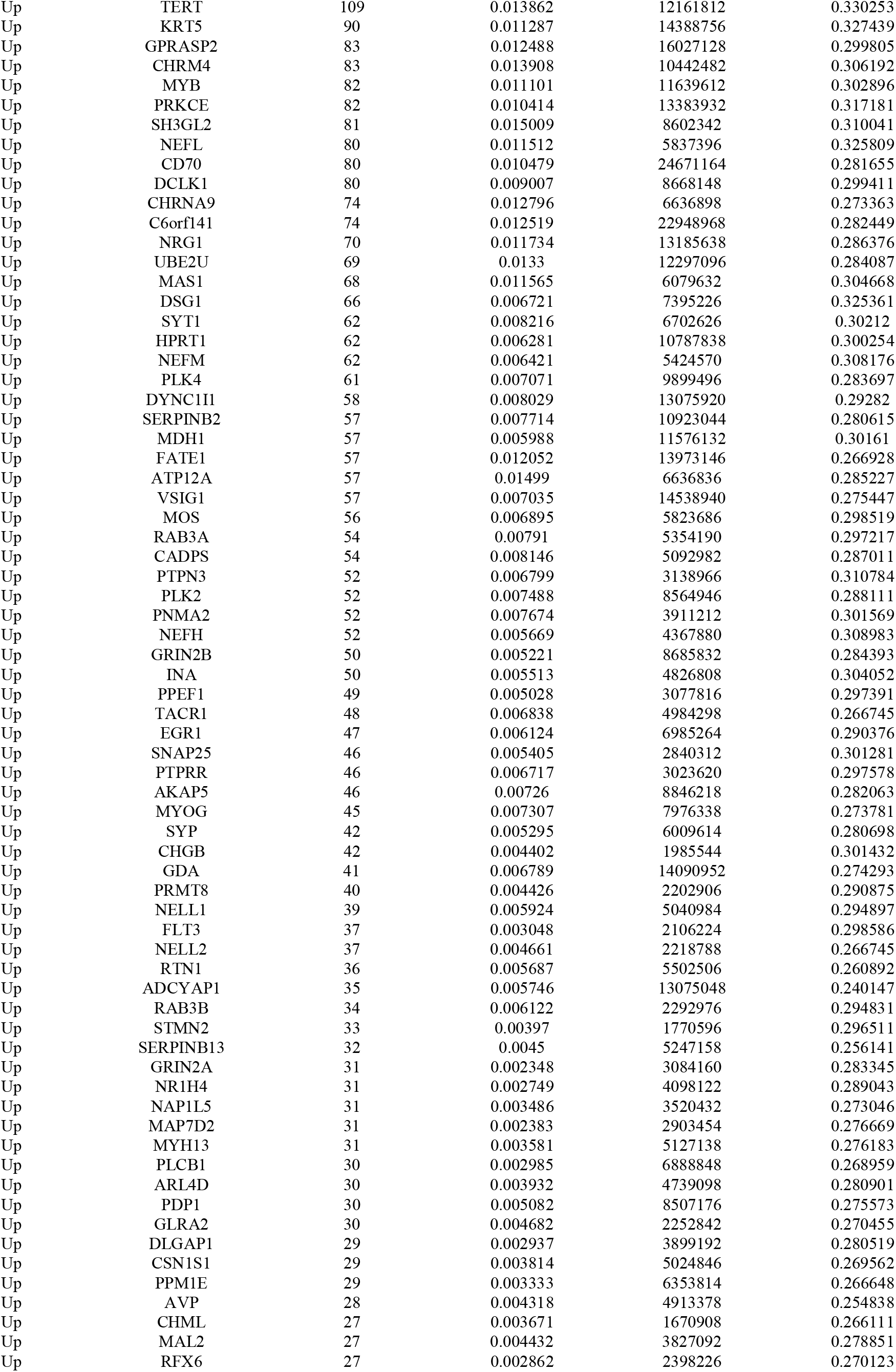
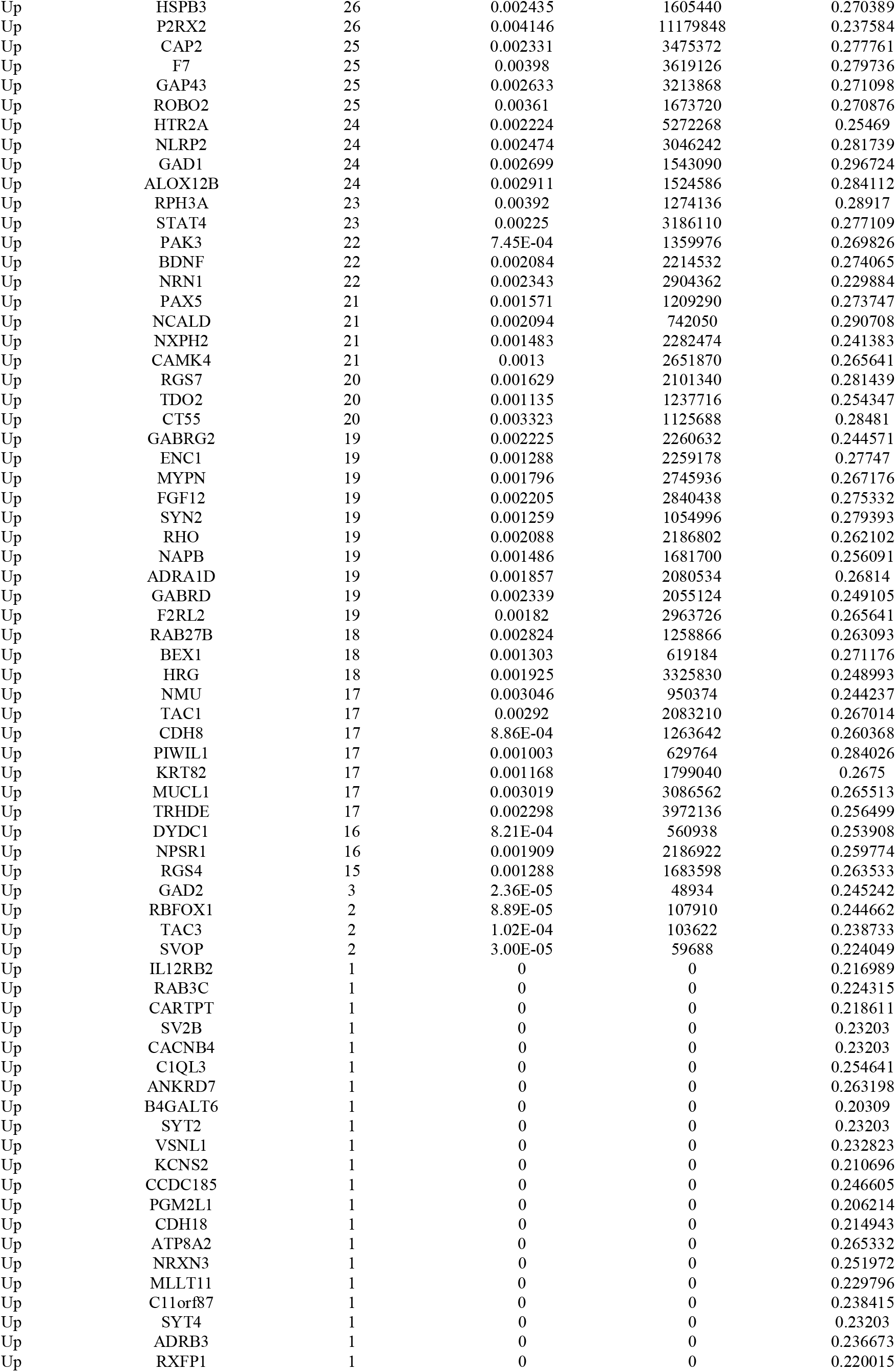
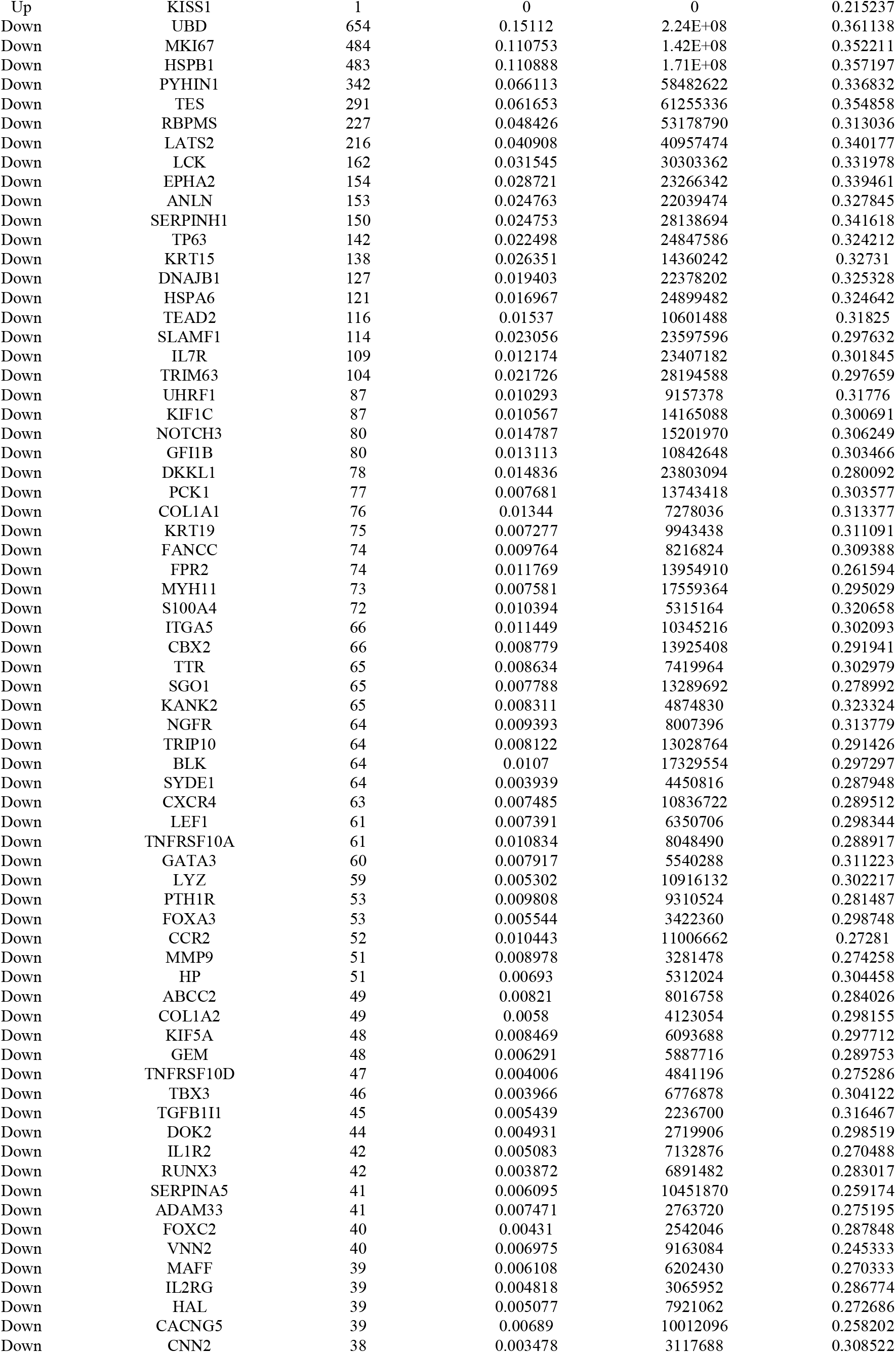
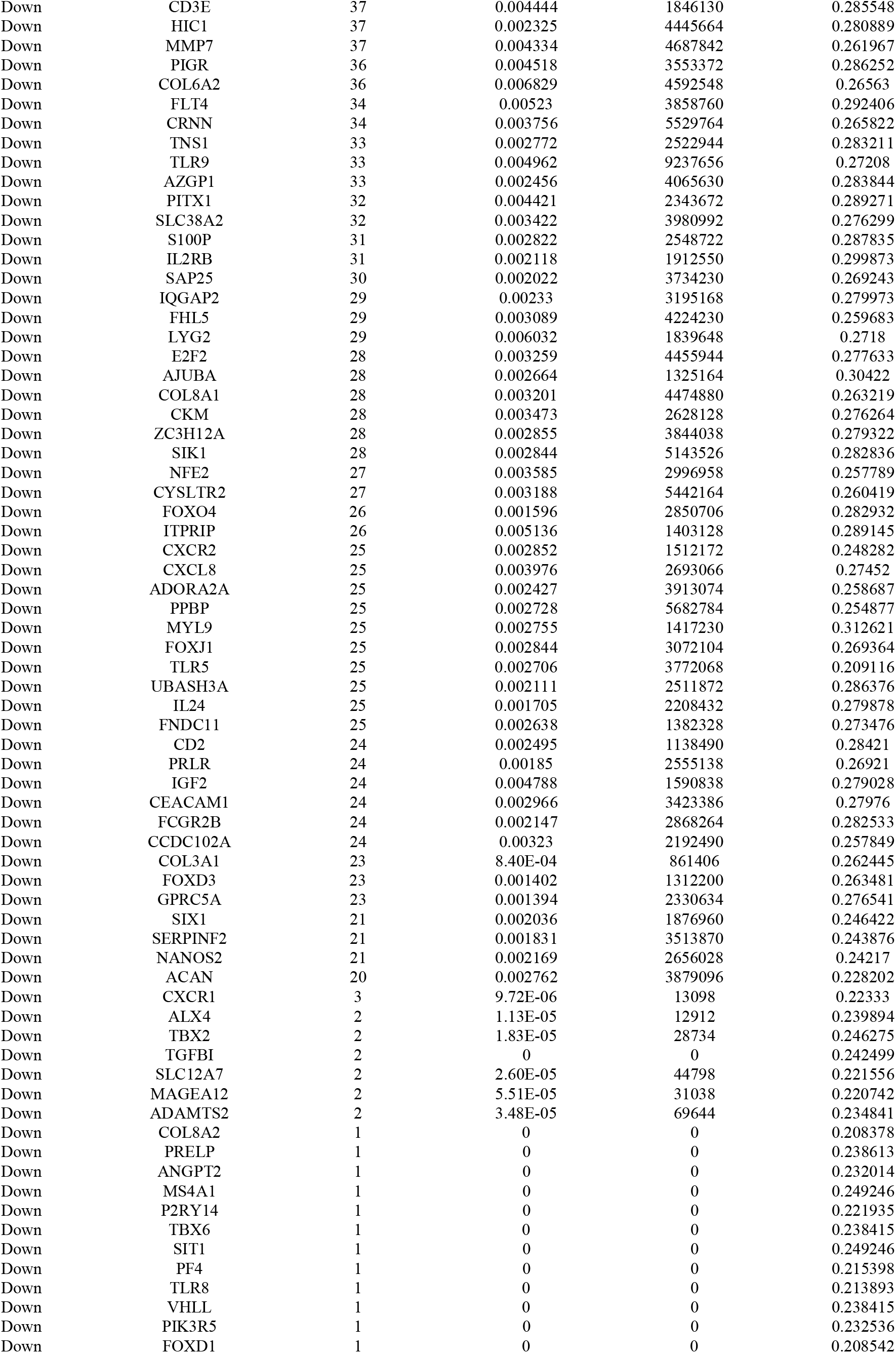
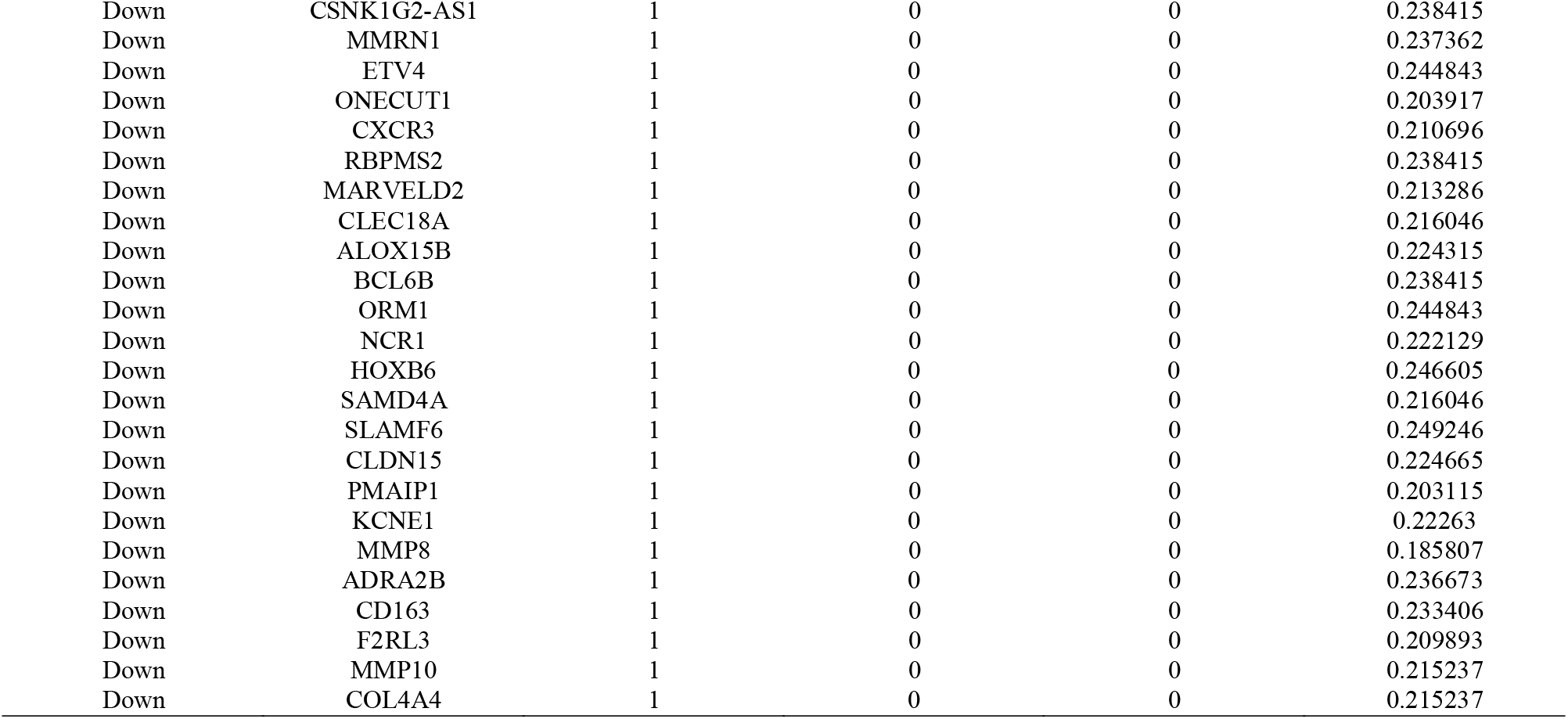
Topology table for up and down regulated genes

**Fig. 4.**
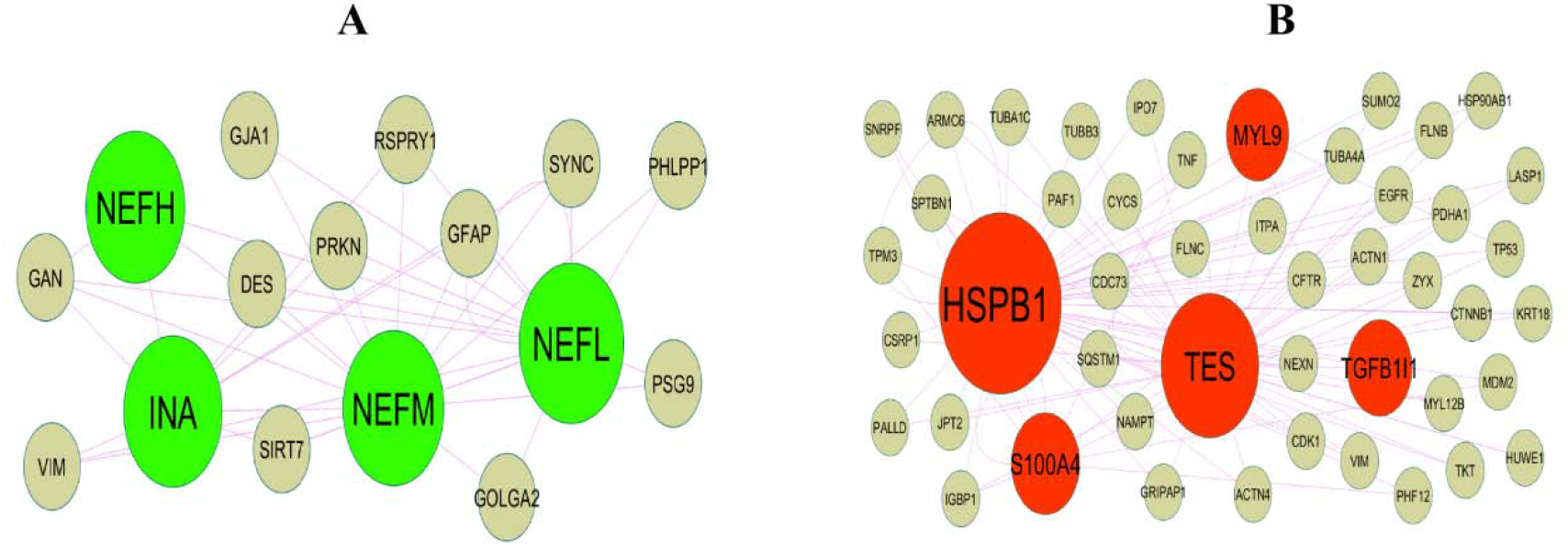
Modules of isolated form PPI of DEGs. (A) The most significant module was obtained from PPI network with 16 nodes and 38 edges for up regulated genes (B) The most significant module was obtained from PPI network with 47 nodes and 96 edges for down regulated genes. Up regulated genes are marked in green; down regulated genes are marked in red.

### MiRNA-hub gene regulatory network construction

As illustrated in Fig. 5, the miRNA-hub gene regulatory network consists of 2316 nodes (301 hub genes and 2015 miRNAs) and 11018 edges. Notably, NEFL targeted 78 miRNAs (ex; hsa-mir-378a-3p); NSF targeted 45 miRNAs (ex; hsa-mir-100-5p); ELAVL2 targeted 43 miRNAs (ex; hsa-mir-27a-3p); PRKCE targeted 42 miRNAs (ex; hsa-mir-513a-5p); MYB targeted 34 miRNAs (ex; hsa-mir-15b-5p); ANLN targeted 126 miRNAs (ex; hsa-mir-138-1-3p); MKI67 targeted 124 miRNAs (ex; hsa-mir-548d-5p); SERPINH1 targeted 83 miRNAs (ex; hsa-mir-4726-3p); LATS2 targeted 70 miRNAs (ex; hsa-mir-320d) and TES targeted 70 miRNAs (ex; hsa-mir-1202) and are listed in Table 5.

**Fig. 5.**
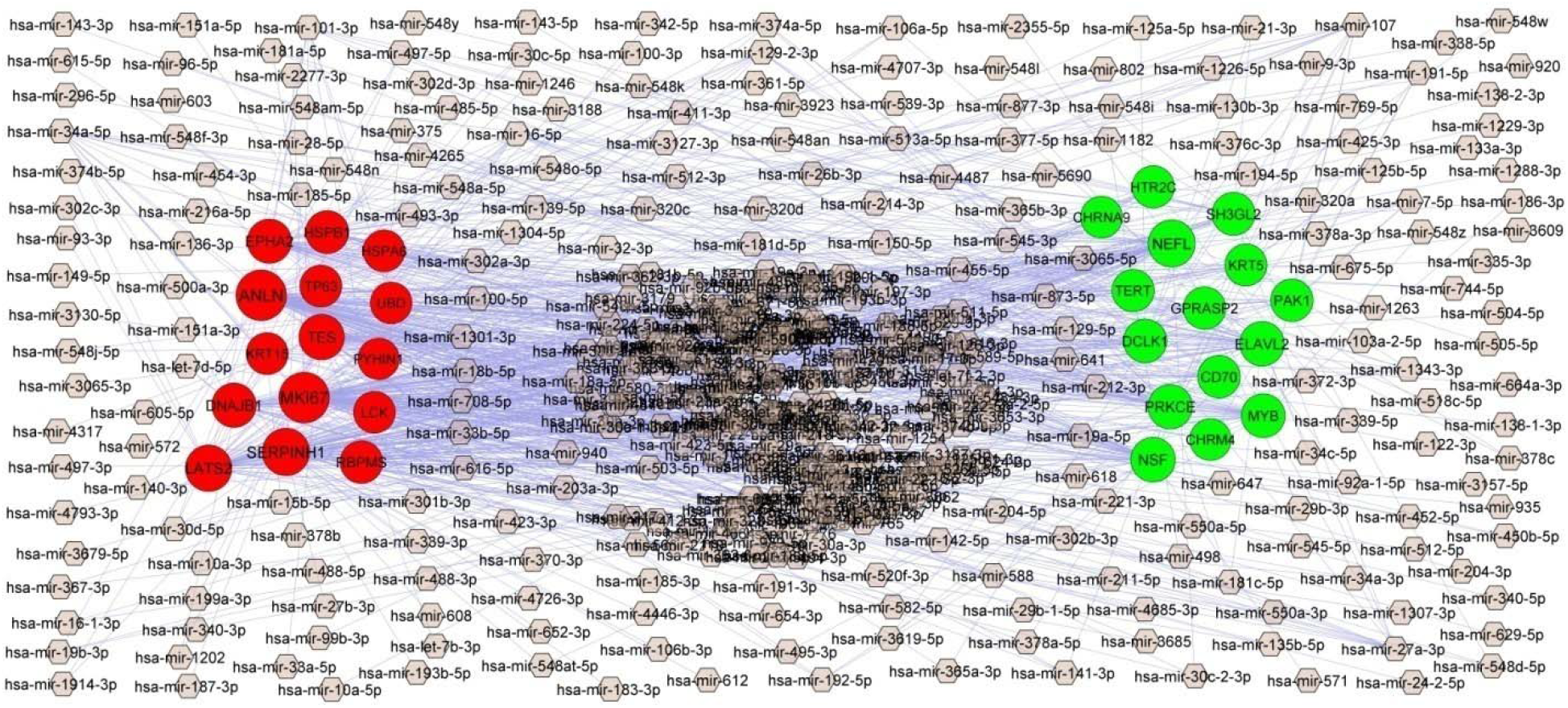
MiRNA - hub gene regulatory network. The purple color diamond nodes represent the key miRNAs; up regulated genes are marked in green; down regulated genes are marked in red.

**Table 5.** miRNA - hub gene and TF – hub gene interaction

| Regulation | Hub Genes | Degree | MicroRNA | Regulation | Hub Genes | Degree | TF |
| --- | --- | --- | --- | --- | --- | --- | --- |
| Up | NEFL | 78 | hsa-mir-378a-3p | Up | ELAVL2 | 15 | NRF1 |
| Up | NSF | 45 | hsa-mir-100-5p | Up | MYB | 15 | STAT1 |
| Up | ELAVL2 | 43 | hsa-mir-27a-3p | Up | DCLK1 | 14 | YY1 |
| Up | PRKCE | 42 | hsa-mir-513a-5p | Up | CD70 | 10 | PPARG |
| Up | MYB | 34 | hsa-mir-15b-5p | Up | TERT | 10 | E2F6 |
| Up | DCLK1 | 28 | hsa-mir-365a-3p | Up | PRKCE | 9 | GATA2 |
| Up | SH3GL2 | 23 | hsa-mir-548o-5p | Up | PAK1 | 7 | REL |
| Up | CD70 | 17 | hsa-mir-374a-5p | Up | KRT5 | 7 | SREBF1 |
| Up | PAK1 | 15 | hsa-mir-107 | Up | NSF | 7 | SRF |
| Up | GPRASP2 | 15 | hsa-mir-518c-5p | Up | HTR2C | 6 | JUND |
| Up | TERT | 13 | hsa-mir-498 | Up | CHRM4 | 6 | PAX2 |
| Up | HTR2C | 12 | hsa-mir-339-5p | Up | CHRNA9 | 5 | JUN |
| Up | KRT5 | 9 | hsa-mir-125b-5p | Up | SH3GL2 | 5 | EN1 |
| Up | CHRNA9 | 6 | hsa-mir-23b-3p | Up | NEFL | 5 | SREBF2 |
| Up | CHRM4 | 4 | hsa-mir-16-5p | Up | GPRASP2 | 2 | HNF4A |
| Down | ANLN | 126 | hsa-mir-138-1-3p | Down | TP63 | 17 | ZNF354C |
| Down | MKI67 | 124 | hsa-mir-548d-5p | Down | TES | 11 | ARID3A |
| Down | SERPINH1 | 83 | hsa-mir-4726-3p | Down | LCK | 10 | FOXL1 |
| Down | LATS2 | 70 | hsa-mir-320d | Down | HSPB1 | 9 | RUNX2 |
| Down | TES | 57 | hsa-mir-1202 | Down | PYHIN1 | 8 | TP53 |
| Down | DNAJB1 | 42 | hsa-mir-191-5p | Down | SERPINH1 | 8 | FOS |
| Down | EPHA2 | 40 | hsa-mir-450b-5p | Down | ANLN | 8 | TFAP2C |
| Down | HSPB1 | 22 | hsa-mir-548f-3p | Down | MKI67 | 7 | FOXC1 |
| Down | RBPMS | 19 | hsa-mir-214-3p | Down | DNAJB1 | 7 | EGR1 |
| Down | TP63 | 9 | hsa-mir-139-5p | Down | UBD | 6 | GATA1 |
| Down | KRT15 | 9 | hsa-mir-1343-3p | Down | HSPA6 | 5 | MEF2A |
| Down | HSPA6 | 8 | hsa-mir-204-3p | Down | LATS2 | 5 | HOXA5 |
| Down | LCK | 6 | hsa-mir-101-3p | Down | EPHA2 | 5 | ELK1 |
| Down | UBD | 5 | hsa-mir-21-3p | Down | RBPMS | 4 | SPIB |
| Down | PYHIN1 | 2 | hsa-mir-375 | Down | KRT15 | 3 | SOX5 |

### TF-hub gene regulatory network construction

As illustrated in Fig. 6, the TF-hub gene regulatory network consists of 386 (305 hub genes, 81 TFs) and 2380 edges. Notably, ELAVL2 targeted 15 TFs (ex; NRF1); MYB targeted 15 TFs (ex; STAT1); DCLK1 targeted 14 TFs (ex; YY1); CD70 targeted 10 TFs (ex; PPARG); TERT targeted 10 TFs (ex; E2F6); TP63 targeted 17 TFs (ex; ZNF354C); TES targeted 11 TFs (ex; ARID3A); LCK targeted 10 TFs (ex; FOXL1); HSPB1 targeted 9 TFs (ex; RUNX2) and PYHIN1 targeted 8 TFs (ex; TP53) and are listed in Table 5.

**Fig. 6.**
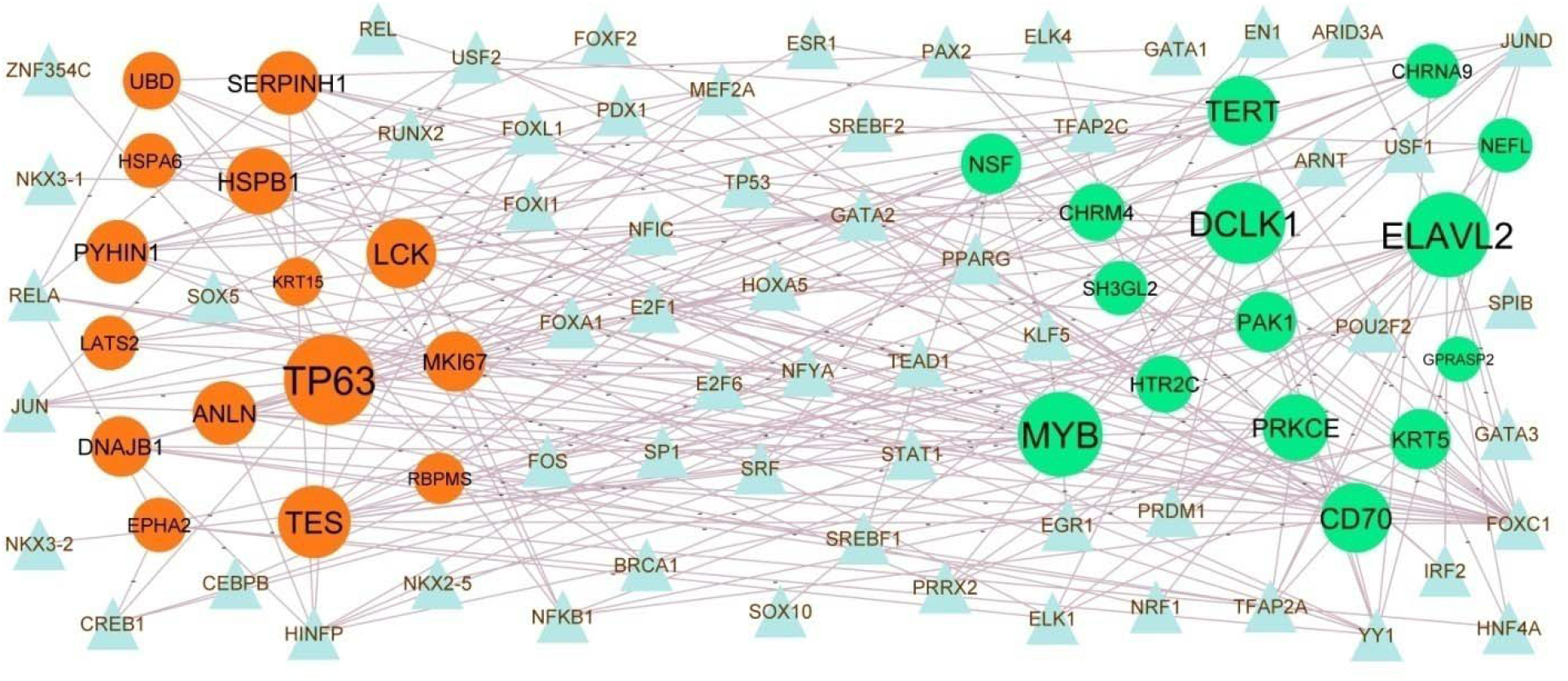
TF - hub gene regulatory network. The blue color triangle nodes represent the key TFs; up regulated genes are marked in green; down regulated genes are marked in red.

### Validation of hub genes by receiver operating characteristic curve (ROC) analysis

First of all, we performed the ROC curve analysis among 10 hub genes. The results showed that PAK1, ELAVL2, NSF, HTR2C, TERT, UBD, MKI67, HSPB1, PYHIN1 and TES achieved an AUC value of□ >□0.9, demonstrating that these eight genes have high sensitivity and specificity for AD, suggesting they can be served as biomarkers for the diagnosis of AD (Fig. 7A-Fig. 7J).

**Fig. 7.**
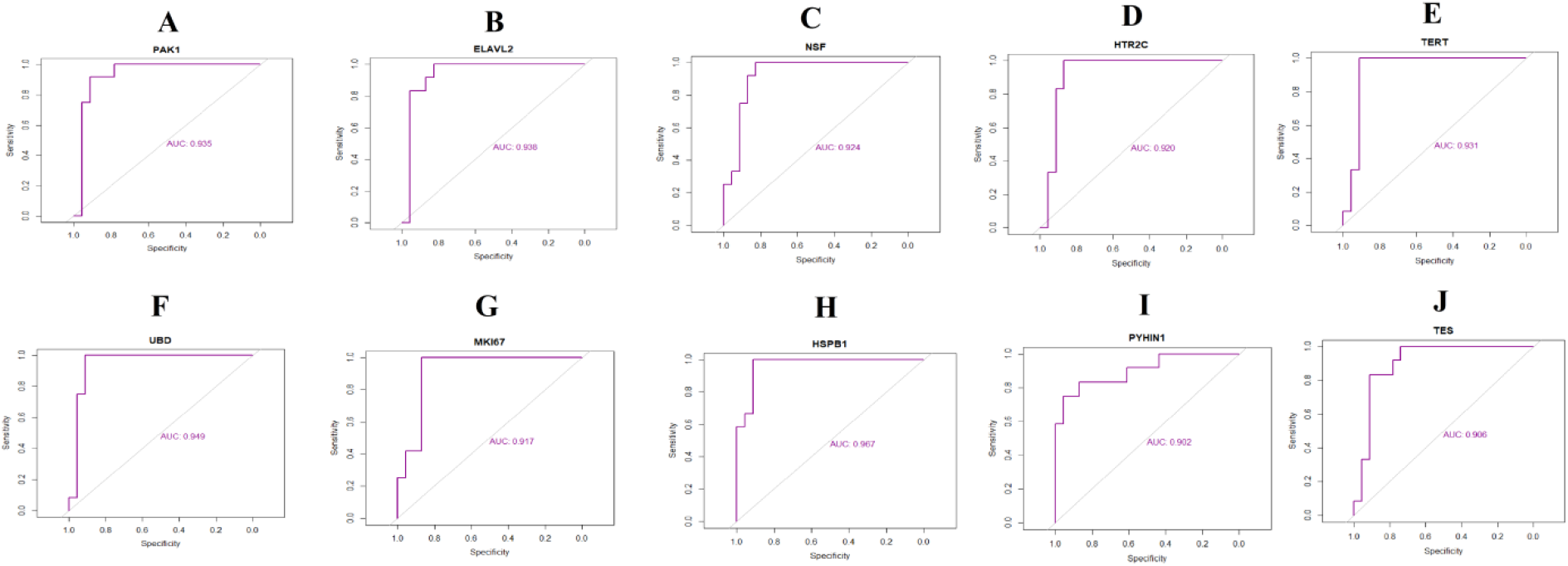
ROC curve validated the sensitivity, specificity of hub genes as a predictive biomarker for Alzheimer’s disease diagnosis. A) PAK1 B) ELAVL2 C) NSF D) HTR2C E) TERT F) UBD G) MKI67 H) HSPB1 I) PYHIN1 J) TES

## Discussion

Genetic modification is widely accepted as a robust molecular event of AD. Unfortunately, little knowledge has focused on genetic modification in AD. In the present study, essential candidate genes and enriched pathways of AD were identified by bioinformatics analysis. We extracted the expression profiling by high throughput sequencing data from GSE125583 and obtained 479 up regulated and 477 down regulated genes in AD. A new study reported the CRH (corticotropin releasing hormone) [32], SST (somatostatin) [33] and GABRR2 [34] could be a biomarkers of AD. TAC1 has been shown to be prognostic biomarkers and therapeutic targets in multiple sclerosis [35], but this gene might be responsible for progression of AD. VGF (VGF nerve growth factor inducible) is a key regulator in patients with amyotrophic lateral sclerosis [36], but this gene might be linked with development of AD. A previous study indicated that the expression of MMP8 [37] and MMP7 [38] are associated with development of dementia.

Functional enrichment analysis was performed based on the GO database and the REACTOME database to screen the key pathways and genes to provide insights into the physiological functions and progress of AD. Neuronal system [39], neurotransmitter receptors and postsynaptic signal transmission [40], signaling by GPCR [41], signal transduction [42], GPCR ligand binding [43], immune system [44], hemostasis [45], cell surface interactions at the vascular wall [46], cell periphery [47], channel activity [48], response to stimulus [49], plasma membrane [50], extracellular region [51], signaling receptor activity [52] and G protein-coupled receptor activity [53] were liable for progression of AD. Matsuzaki et al. [54], Shepard et al. [55], Lennertz et al. [56], Klemettilä et al. [57], Ullah et al. [58], Kim et al. [59], Sasayama et al. [60], Li et al. [61], Kowalczyk et al. [62] and Sun et al. [63] revealed that the expression of ADCYAP1, NPAS4, NPSR1, HTR2C, GABRB2, ALOX12B, ADRB3, EGR3, HSPA1A and IL3RA are associated with progression of schizophrenia, but these genes might be novel target for AD. GABRA6 [64], GABRA1 [65], GABRG2 [66], SLC4A10 [67], HCN1 [68], GABRA4 [69], KCNC2 [70], SCN2A [71], SYN2 [72], FGF12 [73], SCN8A [74], OLFM3 [75], PLCB1 [76], KCNQ5 [77], TUBB2A [78], SIK1 [79], ABCC2 [80], SLC6A12 [81] and COL6A2 [82] have been reported to be closely related to the occurrence and development of epilepsy, but these genes might be novel target for AD. Previous investigation demonstrates that RGS4 [83], CXCL11 [84], EGR1 [85], CALB1 [86], BDNF (brain derived neurotrophic factor) [87], TERT (telomerase reverse transcriptase) [88], NEFL (neurofilament light) [89], SNAP25 [90], RPH3A [91], NRN1 [92], SYT1 [93], GRIN2B [94], AVP (arginine vasopressin) [95], VSNL1 [96], HTR2A [97], PAK3 [98], STXBP5L [99], HCRTR2 [100], SYP (synaptophysin) [101], SYT10 [102], PRKCE (protein kinase C epsilon) [103], NRG1 [104], KISS1 [105], NRXN3 [106], RAB3A [107], IGF1 [108], PLK2 [109], CBLN4 [110], CAP2 [111], SV2B [112], CAMK4 [113], INA (internexin neuronal intermediate filament protein alpha) [114], GAP43 [115], TTR (transthyretin) [116], CXCR2 [117], IL1R2 [118], CXCR4 [119], CCR2 [120], MYOCD (myocardin) [121], S100A12 [122], CXCR3 [123], PROK2 [124], CXCL8 [125], RGS1 [126], NOTCH3 [127], P2RX7 [128], NGFR (nerve growth factor receptor) [129], GDF15 [130], CR1 [131], ADORA2A [132], GPER1 [133], FCGR2B [134], MMP9 [135], CNN2 [136], C5AR1 [137], CCL15 [138], RBPMS (RNA binding protein, mRNA processing factor) [139], TBX2 [140], FPR2 [141], PCK1 [142], ADH1B [143], TLR5 [144], C7 [145] and CD163 [146] are thought to contribute to AD development and it has been reported to act as a potential biomarkers for AD treatment. IL1RAPL2 [147], GAD1 [148], EPHA5 [149], HTR1A [150], CADPS (calcium dependent secretion activator) [151], GABRG3 [152], LRFN5 [153] and CXCR1 [154] are expressed in autism spectrum disorders, but these genes might be novel target for AD. GAD2 [155], TAC3 [156], NCR1 [157], IL7R [158], GATA3 [159], FCRL3 [160] and IL18R1 [161] have been reported as most promising biomarkers for early diagnosis and prognosis prediction of multiple sclerosis, but these genes might be novel target for AD. FLT3 [162], CHRNB4 [163], PAK1 [164], CHRNB3 [165], GRIN2A [166], CHRNA6 [167], PENK (proenkephalin) [168], CHRNB4 [169], IGF2 [170], TLR9 [171] and CACNG5 [172] were revealed to serve an important role in Parkinson’s disease, but these genes might be novel target for AD. The expression of SLC30A3 [173], NEFH (neurofilament heavy) [174], SYNGR4 [175], S100A4 [176], HSPB1 [177] and KIF5A [178] are associated with amyotrophic lateral sclerosis, but these genes might be novel target for AD.

Based on the node degree, betweenness centrality, stress centrality and closeness centrality in PPI network and modules, the hub genes were ranked. These identified hub genes were functioned as a group, and might play a crucial role in AD. ELAVL2 has been identified in autism spectrum disorders [179], but this gene might be novel target for AD. NEFM (neurofilament medium) was revealed to be expressed in AD [180]. NSF (N-ethylmaleimide sensitive factor, vesicle fusing ATPase), UBD (ubiquitin D), MKI67, PYHIN1, TES (testin LIM domain protein), MYL9 and TGFB1I1 might be the novel biomarkers for AD.

By analyzing the miRNA-hub gene regulatory network construction and TF-hub gene regulatory network, a number of hub genes, miRNAs and TFs were identified that might provide novel approaches for therapeutic investigations of AD. CD70 has been shown as a promising biomarker in multiple sclerosis [181], but this gene might be novel target for AD. Hsa-mir-378a-3p [182], hsa-mir-100-5p [182], hsa-mir-27a-3p [183], hsa-mir-15b-5p [184], hsa-mir-320d [185], NRF1 [186], STAT1 [187], YY1 [188], PPARG (peroxisome proliferator-activated receptor gamma) [189] and TP53 [190] have been shown to be an important role in AD. MYB (MYB proto-oncogene, transcription factor), ANLN (anillin actin binding protein), SERPINH1, LATS2, DCLK1, TP63, LCK (LCK proto-oncogene, Src family tyrosine kinase), hsa-mir-513a-5p, hsa-mir-138-1-3p, hsa-mir-548d-5p, hsa-mir-4726-3p, hsa-mir-1202, E2F6, ZNF354C, ARID3A, FOXL1 and RUNX2 might be the novel biomarkers for AD.

In conclusion, integrated bioinformatics analysis of multiple datasets of newly diagnosed AD patients and healthy controls was performed. Common DEGs were identified that are significantly enriched in various pathways, followed by evaluation of the prognostic value for patients with AD. Notably, most of the prognostic genes are neuron and immune related genes. The results of this investigation increase our understanding of the molecular drivers that underlie AD initiation and progression, and the identified key genes and pathways constitute potential therapeutic targets.

## Acknowledgement

I thank Brad A Friedman, Genentech, Inc., Bioinformatics, South San Francisco, CA, USA, very much, the author who deposited their profiling by high throughput sequencing dataset GSE125583, into the public GEO database.

## Ethical approval

This article does not contain any studies with human participants or animals performed by any of the authors.

## Informed consent

No informed consent because this study does not contain human or animals participants.

## Availability of data and materials

The datasets supporting the conclusions of this article are available in the GEO (Gene Expression Omnibus) (https://www.ncbi.nlm.nih.gov/geo/) repository. [(GSE125583) (https://www.ncbi.nlm.nih.gov/geo/query/acc.cgi?acc=GSE125583)]

## Consent for publication

Not applicable.

## Author Contributions

B. V. - Writing original draft, and review and editing

C. V. - Software and investigation

